# Prioritizing conservation actions for Pacific salmon in Canada

**DOI:** 10.1101/2020.02.03.931691

**Authors:** Jessica C. Walsh, Katrina Connors, Eric Hertz, Laura Kehoe, Tara G. Martin, Brendan Connors, Michael J. Bradford, Cameron Freshwater, Alejandro Frid, Jessica Halverson, Jonathan W. Moore, Michael H.H. Price, John D. Reynolds

## Abstract

1. Current investment in conservation is insufficient to adequately protect and recover all ecosystems and species. The challenge of allocating limited funds is acute for Pacific salmon (*Oncorhynchus* spp.) in Canada, which lack a strategic approach to ensure that resources are spent on actions that would cost-effectively recover diminished populations.
2. We applied the Priority Threat Management framework to prioritize strategies that are most likely to maximize the number of thriving Pacific salmon populations on the Central Coast of British Columbia, Canada. These included 79 genetically, ecologically and spatially distinct population groups called Conservation Units (CUs) for five salmon species. This region has high salmon biodiversity and spans the territories of four First Nations: the Heiltsuk, Nuxalk, Kitasoo/Xai’xais and Wuikinuxv.
3. Using structured expert elicitation of Indigenous and other experts, we quantified the estimated benefits, costs and feasibility of implementing 10 strategies. Under a business-as-usual scenario (i.e., no additional investments in salmon conservation or management), experts predicted that only one in four CUs would have >50% chance of achieving a thriving status within 20 years. Limiting future industrial development, which was predicted to safeguard CUs from future declines, was identified as the most cost-effective strategy. Investment in three strategies: 1) removal of artificial barriers to fish migration, 2) watershed protection, and 3) stream restoration – at 11.3M CAD per year – was predicted to result in nearly half (34 of 79) of the CUs having a >60% chance of meeting the conservation objective.
4. If all conservation strategies were implemented, experts estimated a >50% probability of achieving a thriving status for 78 of 79 CUs, at an annual cost of 17.3M CAD. However, even with the implementation of all strategies, most sockeye salmon CUs were unlikely to achieve higher probability targets of reaching the objective.
5. *Policy implications*: We illustrate how Priority Threat Management can incorporate the perspectives and expertise of Indigenous peoples and other experts to evaluate and prioritize conservation strategies based on their cost, benefit and feasibility. Timely investment in the strategies outlined in this assessment could help safeguard and recover Pacific salmon in this region of Canada.

## Introduction

There is an urgent need for strategic planning and prioritization of conservation actions to ensure the persistence and recovery of species and ecosystems, many of which are in decline globally (Martin et al., 2018). However, available resources are inadequate to manage all threats, there is uncertainty around how best to abate these threats, and decision-makers often have limited time and information for prioritizing recovery actions (Martin et al., 2012). Pacific salmon (*Oncorhynchus* spp.) exemplify a group of species that could benefit from strategic planning and prioritization because of their important economic, ecological, and cultural role throughout their range (Mantua et al., 2009). Many populations are diminished or have declined dramatically in recent decades (Governor’s Salmon Recovery Office, 2018; Gustafson et al., 2007; Malick & Cox, 2016; Price, English, Rosenberger, MacDuffee, & Reynolds, 2017). This has led to curtailed fisheries (Ogden et al., 2014; Walters, English, Korman, & Hilborn, 2019), loss of livelihoods, erosion of the cultural identities of Indigenous communities (Garibaldi & Turner, 2004), and potential adverse impacts on coastal ecosystems (Levi et al., 2012). National policies, such as Canada’s Wild Salmon Policy (Fisheries and Oceans Canada, 2005, 2018) recognise the need for a strategic prioritization of management actions (Nelitz, Murray, & Wieckowski, 2008), however, there is currently no strategic framework in Canada for determining how and where to invest limited resources across multiple threats to maximise the probability of recovering salmon populations.

Deciding where and how best to invest in recovery efforts for salmon is a challenge. Factors regulating populations often are complex and poorly understood, but can include stressors such as: overfishing, habitat loss and degradation, barriers to migration, disease, predation, poor survival of juveniles, and climate change (Hoekstra, Bartz, Ruckelshaus, Moslemi, & Harms, 2007; Ruckelshaus, Levin, Johnson, & Kareiva, 2002; Schoen et al., 2017). Furthermore, Pacific salmon have a high degree of local adaptation and policies in the United States and Canada mandate conservation of these evolutionarily distinct units (Waples, 1991). However, existing budgets cannot manage all threats and conserve all salmon populations (Gardner & Pinfold, 2011). There have been many efforts to prioritize potential recovery options for Pacific salmon, such as restoration activities (Beechie, Pess, Roni, & Giannico, 2008; Roni et al., 2002), and guiding recovery plans for at-risk populations (Good, Beechie, McElhany, McClure, & Ruckelshaus, 2007; Kareiva, Marvier, & McClure, 2000). However, it remains difficult to evaluate the costs and benefits of actions across multiple co-occurring populations to ensure that returns on investment are maximized.

Two aspects of the current approach to salmon conservation and recovery planning are problematic. First, the cost-effectiveness of management actions across populations often is not quantified. For example, billions of dollars have been spent on conservation and restoration efforts for Pacific salmon over the last several decades, including enhancement via hatcheries and restoration of spawning and rearing habitats; yet, the effects on populations often are mixed, negligible, or unclear (Barnas, Katz, Hamm, Dias, & Jordan, 2015; Bernhardt et al., 2005). Careful comparison of the cost-effectiveness across management strategies for multiple species and populations could help to maximize salmon recovery potential and protection. Second, resources typically are allocated to the most threatened populations or species (e.g., Cultus Sockeye Recovery Team, 2009; Sakinaw Sockeye Recovery Team, 2005), or those of high public interest (e.g., sockeye or Chinook salmon). Despite the legal and social reasons for prioritizing these diminished populations, they may have the lowest probabilities of recovery and may require the most expensive solutions, compared to less threatened populations with higher recovery potential. If the goal is to increase the total number of healthy populations and their benefits to society, then focusing on the most threatened ones may be suboptimal.

Decision-support tools that explicitly incorporate the cumulative benefits of implementing actions across all species of interest, and the costs and feasibility of these actions, can help identify the most cost-effective management intervention and maximise the benefits to ecosystems and society (Carwardine et al., 2008; Evans et al., 2015). In addition, analyses of the cost-effectiveness that account for complementarity across species can achieve similar conservation outcomes as simple ranking methods but at much lower cost (Chadés et al., 2015). The Priority Threat Management (PTM) framework (Carwardine et al., 2018, 2012; Chadés et al., 2015) accounts for the costs, benefits, feasibility, and complementarity of actions. This framework uses structured expert elicitation to evaluate the cost-effectiveness of strategies that mitigate threats to biodiversity, and identify which strategies would achieve the greatest conservation outcomes across multiple taxa for any given budget. The PTM framework has been applied extensively in Australia (e.g. Carwardine et al., 2012; Chadés et al., 2015) and Canada (Martin et al., 2018). Additional benefits of the PTM framework include greater clarity of objectives, incorporation of Indigenous values and priorities, stakeholder engagement, data centralization and an emphasis on developing baselines necessary to evaluate management performance (Carwardine et al., 2018). PTM uses scientific information, local and traditional knowledge, and expert elicitation to identify where and how to restore imperiled populations. Thus, there is an opportunity to apply PTM to guide systematic planning efforts for Pacific salmon.

We applied the PTM framework to a genus of exceptional economic and cultural importance: Pacific salmon, and in particular, salmon of the Central Coast of British Columbia, Canada. Past PTM exercises have focused on species of conservation concern as their unit of interest. Instead, we focused on populations within and across species, given the importance of genetically-distinct salmon populations, and included all levels of threat status (rather than considering only threatened populations). Our project was a collaboration between academics, NGO staff, federal scientists, the Heiltsuk, Nuxalk, Kitasoo/Xai’xais, and Wuikinuxv Nations - who have lived in the region since time immemorial (Campbell & Butler, 2010; Cannon & Yang, 2006), and the Central Coast Indigenous Resource Alliance (CCIRA – the non-profit society these Nations created for technical support in marine resource management). Our case study, therefore, incorporates the expertise and perspectives of Indigenous people, which are essential from both ecological and social standpoints (Ban et al., 2018), into the development and evaluation of conservation strategies for the cost-effective recovery of Pacific salmon.

## Materials and methods

### Study area

We focused on the Central Coast of BC, including the traditional territories of the four First Nations involved (Figs. 1, S1 & S2, Marine Planning Partnership Initiative, 2015). This region is in temperate rainforest, with relatively low levels of industrial development, supporting hundreds of wild salmon spawning locations that can be delineated into 79 geographically, ecologically, and genetically unique populations of five species: chum (*O. keta*), coho (*O. kisutch*), chinook (*O. tshawytscha*), pink (*O. gorbuscha*) and sockeye (*O. nerka*) salmon. These groups are called Conservation Units (CUs) under Canada’s Wild Salmon Policy (WSP, Fisheries and Oceans Canada, 2005). Four CUs in this region are in the red status zone (at or below their lower biological benchmarks; Holt, Cass, Holtby, & Riddell, 2009a), and 22 CUs are in the amber zone (between the lower and upper biological benchmarks), indicating the need for increased management intervention (Table 1, Connors et al. 2018, Pacific Salmon Explorer, http://www.salmonexplorer.ca/). Twelve CUs are in the green status zone, which suggests relatively high recent spawning abundances compared to benchmarks (Table 1). Over half of the CUs on the Central Coast have insufficient data to assess their biological status (Table 1).

**Figure 1:**
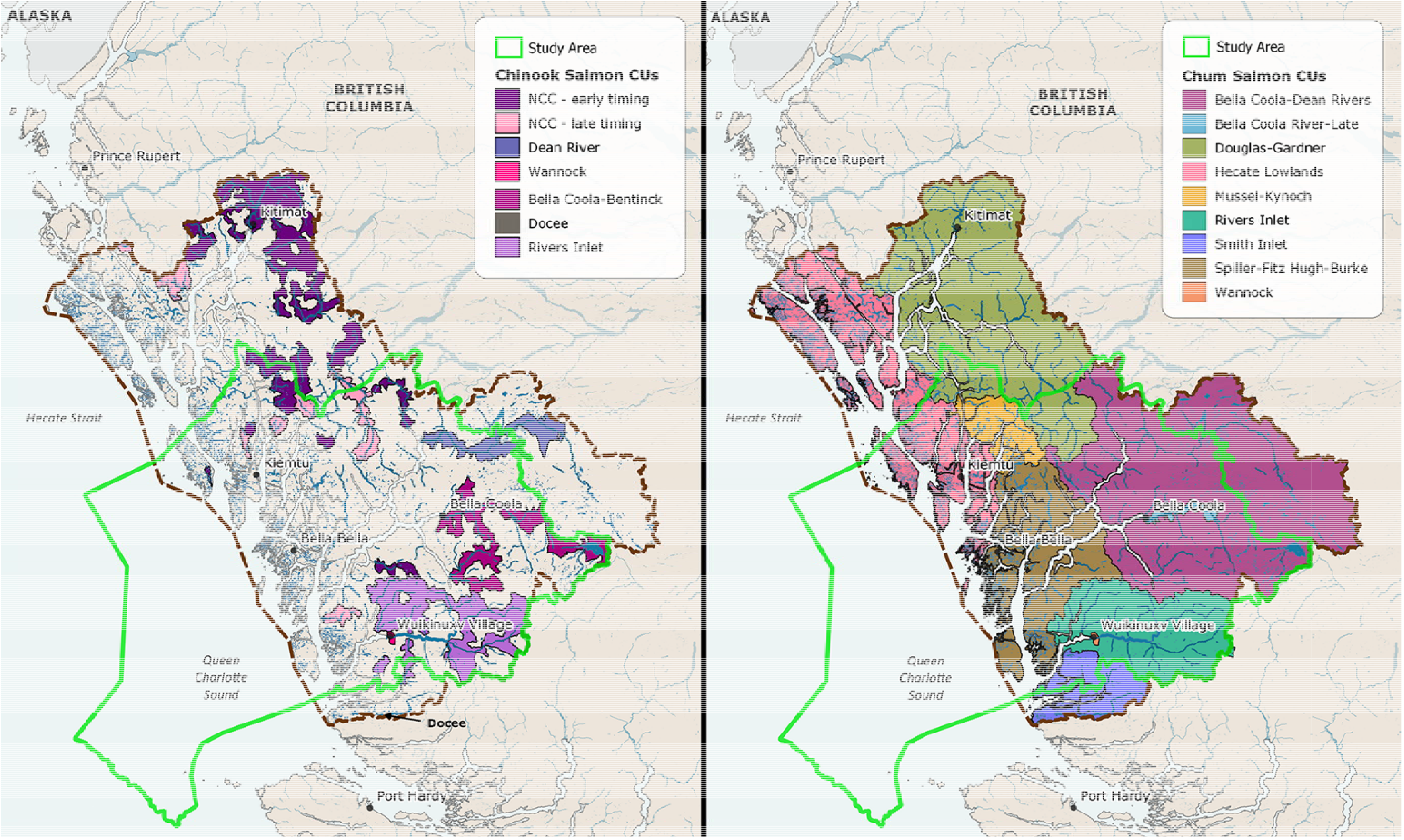
Conservation Units (CUs) for Chinook and chum salmon within the Central Coast of British Columbia, Canada. The study region is outlined in green, showing the combined traditional use territories of the Heiltsuk, Kitasoo/Xai’xais, Nuxalk and Wuikinuxv Nations (total area = 55,266 km^2^). Due to space limitations, maps of coho, pink, river-type sockeye and lake-type sockeye salmon CUs are in Appendix S3: Figs. S1 & S2.

**Table 1:**
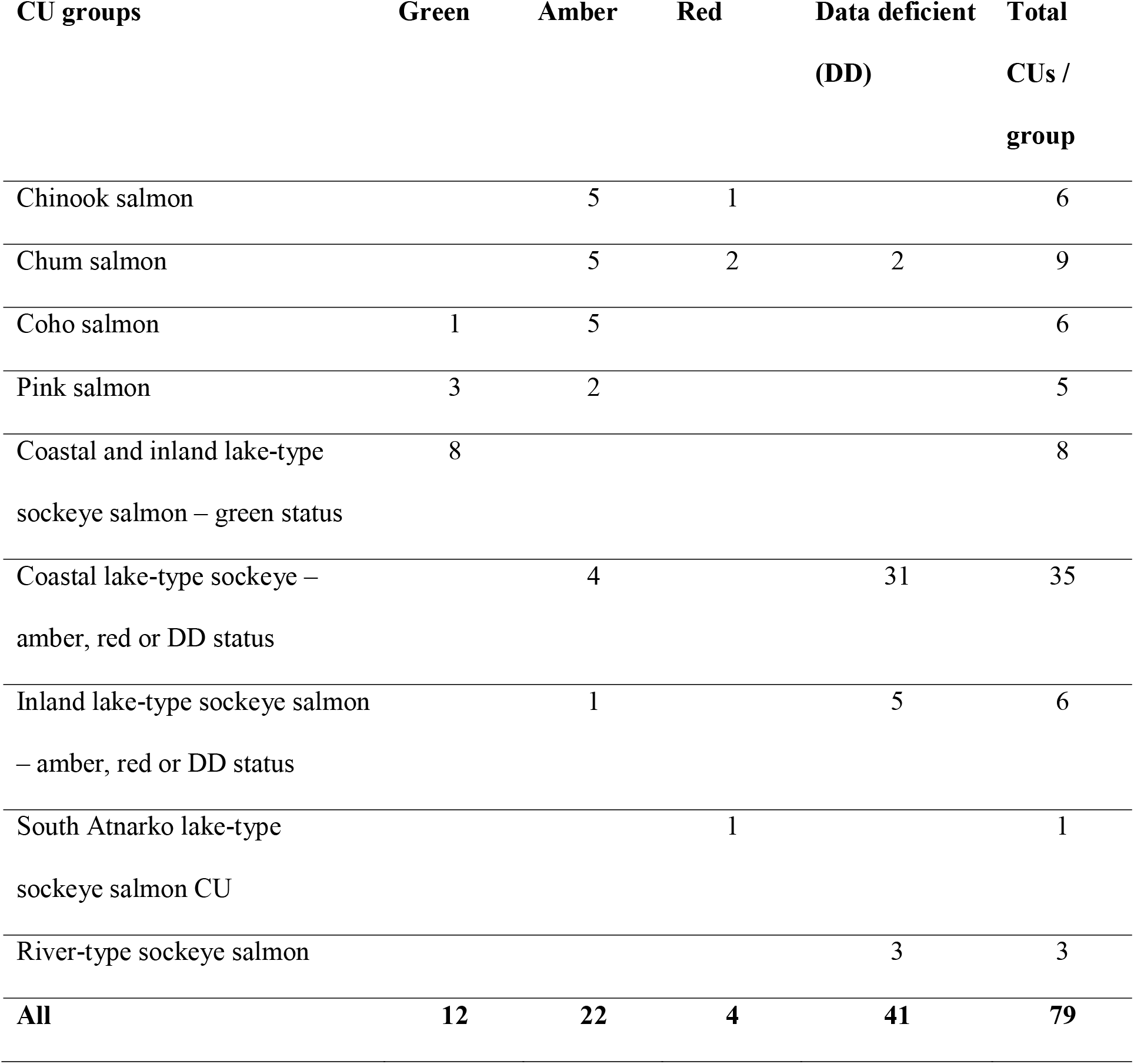
Pacific salmon species and the biological status of their Conservation Units (CUs) on British Columbia’s Central Coast (Fig. 1), which were grouped into 9 CU groups for the analysis. Sockeye CUs were divided into CU groups based on their ecotype, distribution, and biological status (Connors et al., 2018). Status presented here is based on historic spawner abundance benchmarks.

| **CU groups** | **Green** | **Amber** | **Red** | **Data deficient**<br>**(DD)** | **Total**<br>**CUs /**<br>**group** |
| --- | --- | --- | --- | --- | --- |
| Chinook salmon |  | 5 | 1 |  | 6 |
| Chum salmon |  | 5 | 2 | 2 | 9 |
| Coho salmon | 1 | 5 |  |  | 6 |
| Pink salmon | 3 | 2 |  |  | 5 |
| Coastal and inland lake-type<br>sockeye salmon – green status | 8 |  |  |  | 8 |
| Coastal lake-type sockeye –<br>amber, red or DD status |  | 4 |  | 31 | 35 |
| Inland lake-type sockeye salmon<br>– amber, red or DD status |  | 1 |  | 5 | 6 |
| South Atnarko lake-type<br>sockeye salmon CU |  |  | 1 |  | 1 |
| River-type sockeye salmon |  |  |  | 3 | 3 |
| **All** | **12** | **22** | **4** | **41** | **79** |

### Priority threat management framework

The PTM framework (Carwardine et al., 2018, 2012) is an eight-step process that quantifies the cost-effectiveness of management strategies for meeting a stated biodiversity objective, by identifying strategies that abate threats to the biodiversity features of interest (in this case salmon CUs), and estimating the costs, feasibility, and benefits of each strategy. We implemented the PTM framework during consultations with First Nation representatives and three days of structured expert elicitation workshops (Fig. 2). Our research was approved by the Simon Fraser University Office of Research Ethics (permit 2018s0206). The major steps we followed are described below, with further details in Appendix S1.

**Figure 2:**
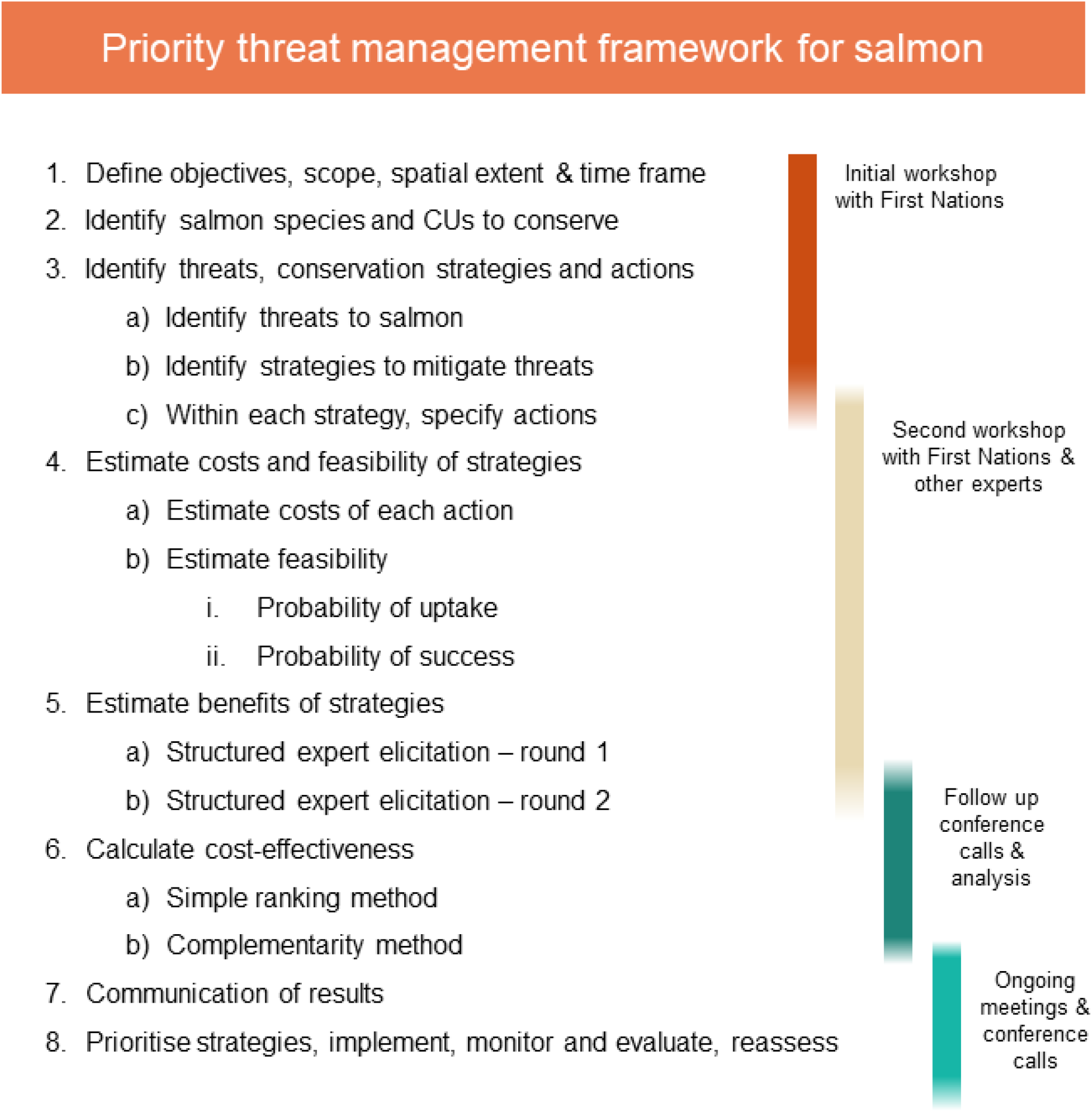
Eight steps of the Priority Threat Management framework. Figure adapted from Carwardine et al. (2019).

#### 1. Define objective, scope and timeframe

The objective was to maximize the number of thriving Pacific salmon CUs on BC’s Central Coast over the next 20 years. We defined a *thriving* CU as fulfilling its ecological function and role, and providing livelihood opportunities for present and future generations. In the context of the Wild Salmon Policy, the objective is analogous to maximizing the number of CUs in the green status zone, which reduces the need for conservation intervention and allows for fishing opportunities, including for First Nation Food, Social, and Ceremonial (FSC) purposes, and commercial and recreational sectors (Fisheries and Oceans Canada, 2005). This objective was defined by the First Nations and experts during the workshops.

#### 2. Identify salmon CUs to conserve

The ‘biodiversity features’ we included in our analysis were all 79 salmon CUs that overlap with our study area. Including CUs with green, amber, red and data deficient (DD) status ensures that conservation strategies (Step 3) are designed to promote recovery of amber and red CUs while avoiding future declines of green status CUs. We grouped the 79 CUs into ‘CU groups’ by species (Table 1). Due to the large number of sockeye salmon CUs, the sockeye CUs were further divided into five life-history groups based on their ecotype (river-type or lake-type), geographic location (inland or coastal) and current level of threat (red/amber/DD status or green status). We assumed that each CU group experienced similar threats and would respond similarly to recovery actions within the region (Table 1). Grouping CUs made the exercise feasible by reducing the workload of experts when estimating the benefits of strategies (Step 5).

#### 3. Identify threats, strategies and actions

We identified threats based on previous summaries of human and environmental pressures in the region (Connors et al., 2018), and a literature review of broader threats to salmon, which were refined by experts at workshops, conference calls, and meetings. These threats included overfishing, habitat loss and degradation due to logging, anthropogenic barriers to freshwater migration, future industrial development in salmon habitats, salmon aquaculture, hatcheries (due to competition and genetic introgression), and predation by marine mammals and other predators. The effects of climate change in marine and freshwater environments were beyond the scope of the strategies we evaluated.

We developed an inventory of strategies, each consisting of several underlying actions (Appendix S2) as well as three combinations of strategies, as experts thought their combined benefits could be synergistic (Table 2). We also developed an enabling strategy that included increased monitoring and assessment of salmon CUs, which experts considered essential for effective implementation and evaluation of all other strategies. This enabling strategy was not included in the cost-effectiveness analysis, because it would not directly result in recovery, but it was assumed that it would be implemented in addition to any other strategy.

**Table 2:**
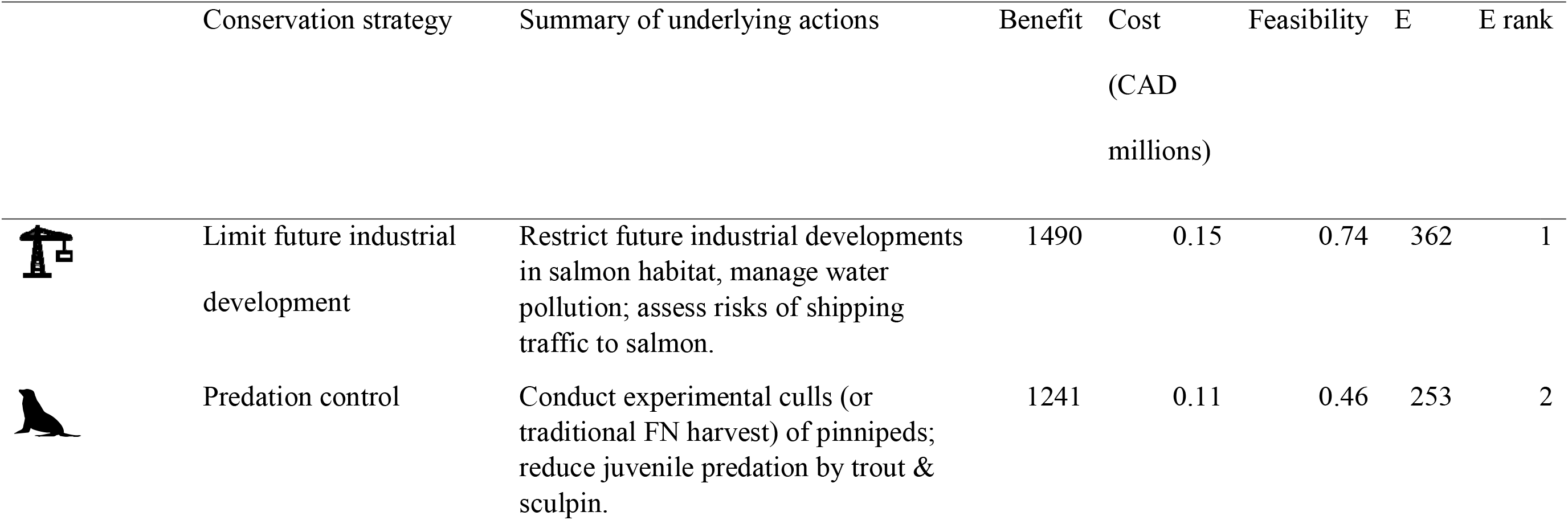
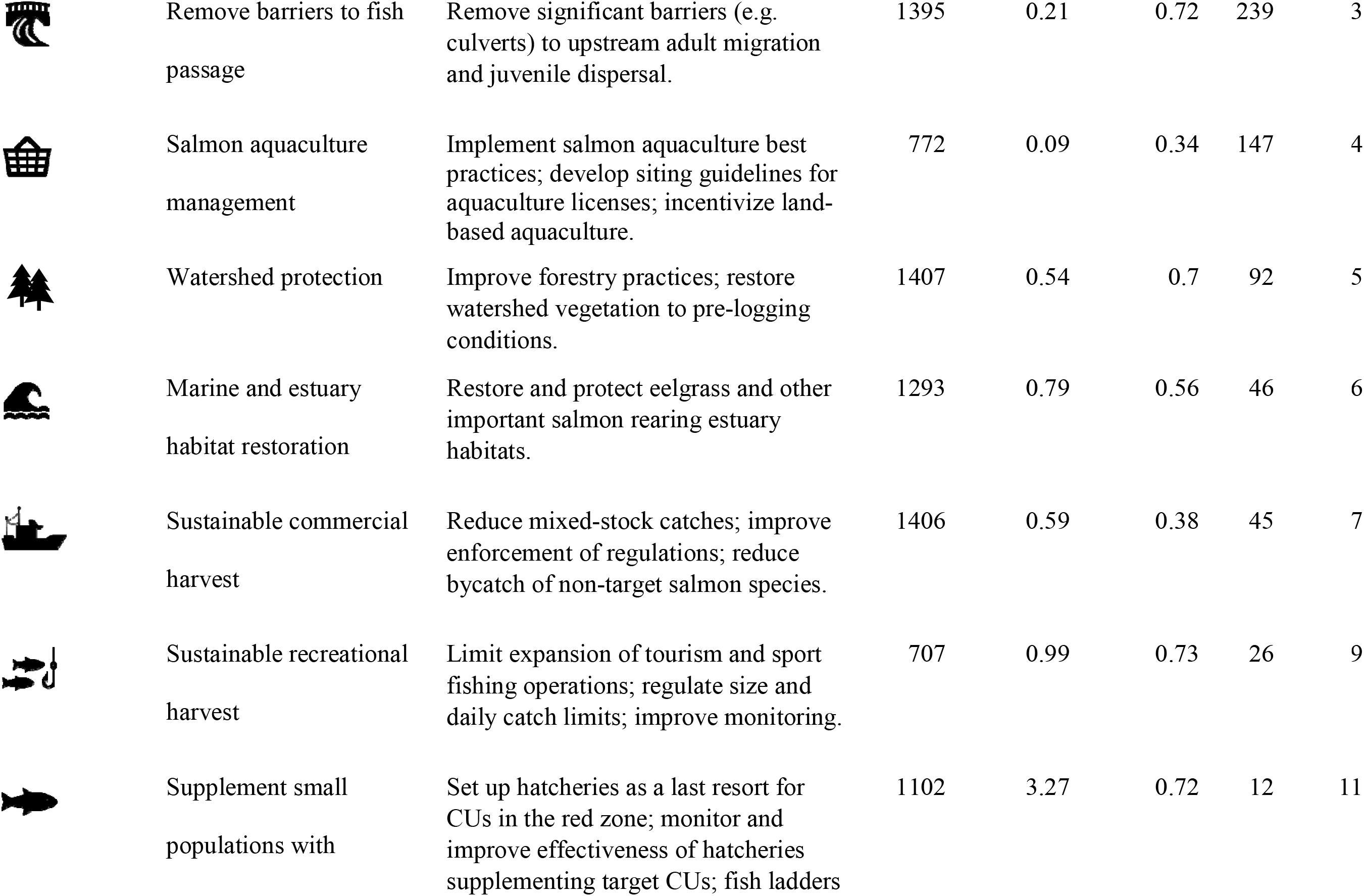
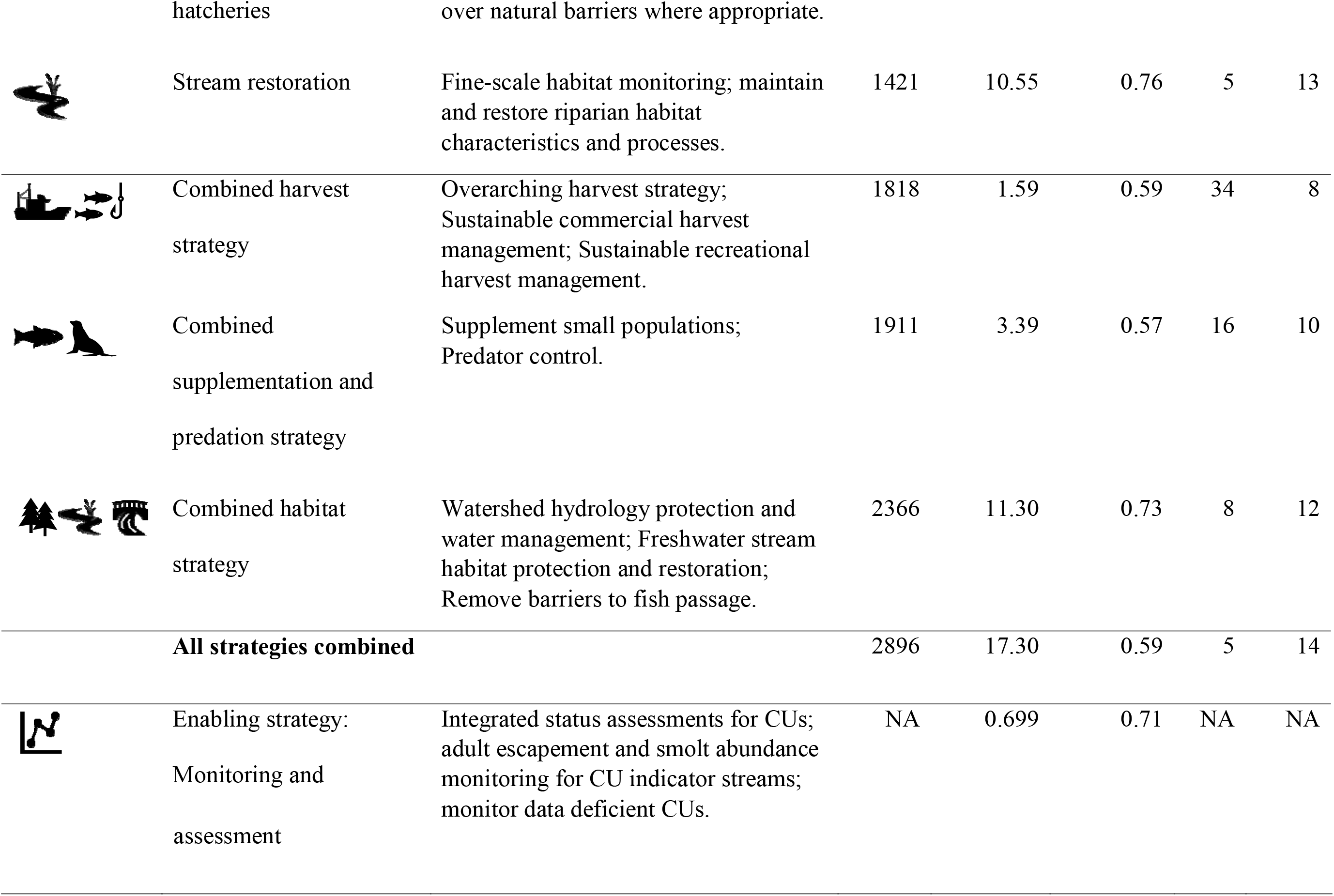
Benefits, costs, feasibility and cost-effectiveness (E) of the ability of each strategy to conserve salmon on British Columbia’s Central Coast. Strategies are ordered by cost-effectiveness from the ranking method. Detailed actions for each strategy are described in Appendix S2. Benefit is the difference between the probability of achieving the conservation objective (i.e., thriving CUs with green status) after 20 years under each strategy and the baseline scenario per CU group, averaged across experts, and summed across all CU groups (Eqn. 1). Cost is the average annual present value divided across the 20-year time frame. Feasibility is the probability of uptake multiplied by the probability of success.

#### 4. Estimate the costs and feasibility of actions

The costs and feasibility of each action within each strategy were quantified during the workshop and finalized in follow-up communication (Appendix S1). The cost of labor, consumables and equipment, capital assets, overheads, monitoring and coordination were estimated for each action, drawing on salmon management reports and the experts’ experience (Iacona et al., 2018). All costs are presented in Canadian dollars (CAD). The feasibility of each strategy was quantified as the product of the average probability of uptake and the average probability of success, across each action.

#### 5. Identify benefits of strategies

We used a structured expert elicitation approach (Hemming, Burgman, Hanea, McBride, & Wintle, 2018; Martin et al., 2012) to elicit the benefits of each strategy for the nine groups of salmon CUs. Fourteen experts (*k*) individually estimated the likely response of each strategy (*i*) for each group of CUs (*j*), defined as the probability that on average each CU within the group would be thriving after 20 years (*P_ijk_*, Fig. S3), along with upper and lower limits and estimates of confidence on a scale of 0-100. Experts also estimated the probability of each CU group achieving this objective in 20 years if no additional management strategies were implemented, which served as a business-as-usual baseline scenario (*P_0jk_*). Experts were asked to consider the effects of climate change and other emerging threats on freshwater and marine habitats, and existing and ongoing management in this baseline scenario. Experts agreed that it was biologically conservative to assume data-deficient CUs are currently in the red status zone (i.e., of conservation concern and below their lower biological benchmark) to ensure estimates of future scenarios were precautionary. Further details on how benefits were estimated are described in Appendix S1.

Following Carwardine et al. (2018), the benefit (*B*_*ij*_) of a strategy for a given CU group was calculated as the change in the probability of achieving the objective under the baseline scenario, compared with the estimated benefit if the strategy was implemented, averaged across experts (*M*_*ij*_ is the number of experts who made predictions for that CU-strategy combination).

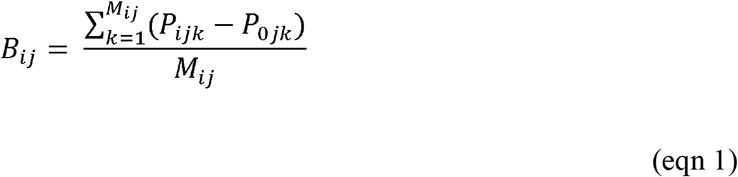

#### 6. Cost-effectiveness calculation

First, using the simple ranking method, we ranked strategies based on their cost-effectiveness (*E_i_*) (Carwardine et al., 2012), using the sum of the cumulative benefits, multiplied by the feasibility per strategy (*F_i_*) divided by the costs per strategy (*C*_*i*_):

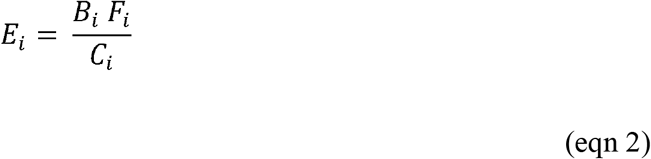

where the total benefit per strategy (*B_i_*) is the sum of the benefits across all CU groups, weighted by the number of CUs occurring in that group (*N_j_*):

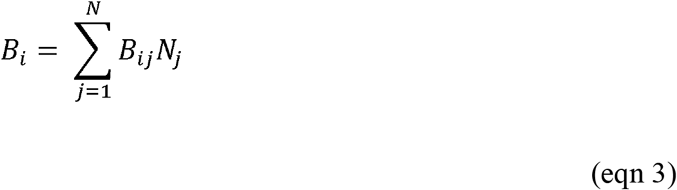

We then used the complementarity approach to identify strategies and their combinations that are predicted to maximize the probability of achieving as many thriving salmon CUs as possible for a given budget (Chadés et al., 2015). For example, rather than choosing two cost-effective strategies that benefit the same CU group, the analysis would select one of those strategies, and another slightly more expensive strategy that benefits a different CU group, thus increasing the number of CUs conserved. To assess complementarity, arbitrary thresholds of ‘conservation’ must be defined to determine the number of CUs that have reached a satisfactory level of protection under any given budget. Here, we selected conservation thresholds of 50%, 60%, and 70% probability of achieving the objective – i.e. being a thriving CU with green status in 20 years.

Uncertainty in the experts’ assessments were assessed by conducting sensitivity analyses to test if the cost-effectiveness results from the simple ranking and complementarity methods were robust to upper and lower estimates of the costs and benefits of each strategy (Appendix S1).

## Results

### Predicted outcomes if no or all strategies were implemented

Under a business-as-usual scenario (i.e., the baseline scenario with no additional investment), only one quarter of salmon CUs on BC’s Central Coast were predicted to have a greater than 50% chance of achieving the conservation objective within the next 20 years (i.e., 6 coho, 5 pink, and 8 ‘green-status lake-type sockeye’ salmon CUs of all 79 CUs, Appendix S3: Table S1, Fig. S3). No CUs were predicted to have a greater than 70% probability of reaching this conservation objective under business-as-usual (Table S1, Fig. 3, Fig. S3). Without additional conservation strategies, declines were predicted for the CUs that currently have green status, such as the 8 CUs in the ‘green-status lake-type’ sockeye salmon group, which were estimated to have 62% probability of thriving in 20 years (Table S1). There was considerable uncertainty in the experts’ best guess estimates of the probability of these CUs still thriving in 20 years (Fig. S3). The lower estimates of expert predictions suggested that under the business-as-usual scenario, no CUs would have a >50% probability of thriving in 20 years and the upper estimates suggest that 19 CUs would have >70% probability of thriving.

**Figure 3:**
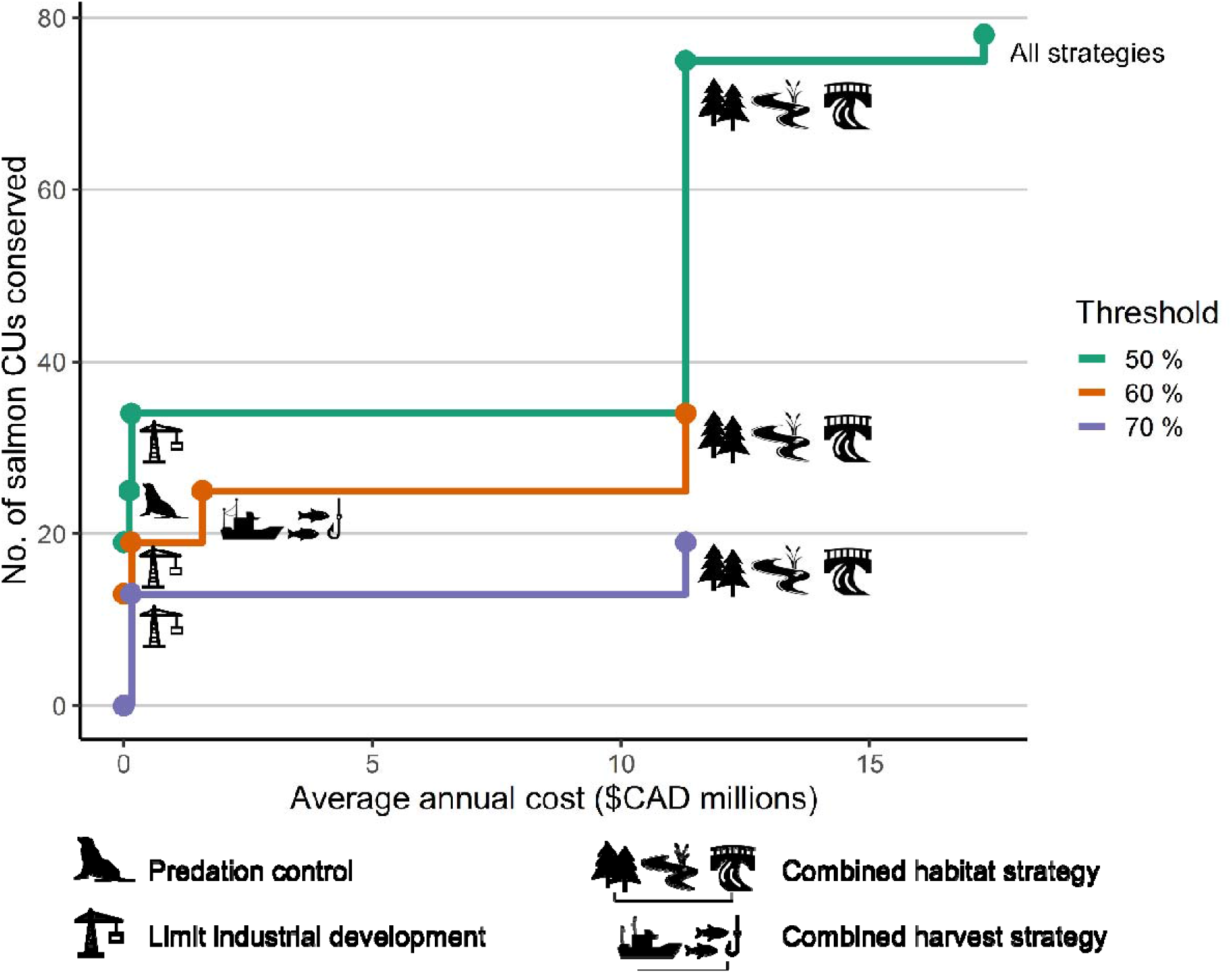
The number of Pacific salmon Conservation Units that would exceed three threshold probabilities (>50%, >60%, & >70%) at different levels of annual investment if complementary sets of strategies were implemented (total = 79 CUs). For example, with an annual budget of $11.3M, the ‘Combined Habitat Strategy’ (i.e., Watershed Protection, Stream Restoration, and Removal of Barriers to Fish Passage) could be implemented, resulting in 75 CUs with >50% probability of success, 34 CUs with >60% probability of success, and 19 CUs with >70% probability of success (shown in Table S1).

Conversely, if all 10 strategies were implemented, all CUs (except the South Atnarko Lake Sockeye CU) were predicted to have >50% probability of thriving in 20 years (Fig. 3). However, only 43% of all CUs – including 6 Chinook, 9 chum, 6 coho, 5 pink, and 8 ‘green-status lake-type sockeye’ salmon – were estimated to reach >60% probability of thriving (Table S1). Amber, red, or data-deficient status sockeye CUs were predicted to have less than 60% probability of meeting the conservation objective even with the implementation of all strategies (Table S1). The experts’ lower estimates suggested that with all 10 strategies, only 5 pink and 8 ‘green-status lake-type sockeye’ salmon CUs would reach >50% probability of achieving thriving status, while the upper estimates suggest that all CUs would reach the >60% threshold, and 34 CUs would have >70% probability of reaching the conservation objective.

The average annual equivalent cost of carrying out all strategies was predicted to be $17.3M per year (min. and max. estimates = $16.1 - $41.4M/year; Table 2). To support the successful implementation of strategies, an additional $0.7M/year would be required to conduct the necessary monitoring and assessment of biological status for all salmon CUs (i.e., enabling strategy in Table 2).

### Cost-effectiveness of strategies

Not all strategies performed equally when comparing their expected benefit (benefit × feasibility) per dollar spent. The most cost-effective strategy predicted to conserve salmon CUs was to ‘Limit Future Industrial Development’ (Table 2, Fig. S4). This strategy aimed to restrict industrial development (e.g. wind farms, aquaculture, and oil and gas infrastructure) in critical areas of salmon spawning and rearing habitat by developing agreements between First Nations and proponents, and maintaining existing tenures. This strategy had the highest benefit to species, and a relatively low cost of $150,000/year for the next 20 years (compared to the average annual cost of individual strategies = $1.73M). On average across all CU groups, ‘Limiting Future Industrial Development’ would result in an increase of 12.6% (SD = 2.6%) probability of achieving a thriving CU compared to the baseline scenario (Table S1). The second most cost-effective strategy was predicted to be ‘Predation control’ (i.e., predator culls which may include – but not be limited to – revitalizing traditional First Nation harvests for problem pinnipeds and trapping of predatory fishes that consume juvenile salmon; Table 2, Fig. S4), which also had a relatively low annual cost of $0.11M.

The ‘Combined Habitat Strategy’ – including ‘Watershed Protection’, ‘Stream Restoration’, and ‘Removal of Barriers to Fish Passage’ – was predicted to be highly beneficial, but had a low cost-effectiveness rank due to higher costs of implementation relative to other strategies (Table 2, Figs. S4 & S5). ‘Stream Restoration’ and ‘Supplement Small Populations’ with hatcheries and other means of enhancement were considered the least cost-effective strategies due primarily to their high costs (Table 2, Fig. S4).

### Complementary strategies to maximize conservation success

We conducted a complementarity analysis to identify which strategies would achieve recovery (or maintain thriving status) for the largest number of CUs under various budgets. Two clear investment thresholds emerge that are consistent across all conservation objective thresholds (Fig. 3, Table S1). The first investment threshold is $0.15M/year, to fund the most cost-effective strategy overall ‘Limit Future Industrial Development’. This strategy was predicted to lead to 19 CUs reaching >60% probability of thriving, and 13 CUs having >70% probability of thriving. Next, for a budget of $11.3M, investing in the ‘Combined Habitat Strategy’ was predicted to deliver the highest number of CU’s conserved across all objective thresholds (34 CUs would reach >60% probability of thriving, and 19 CUs would have >70% probability of thriving). Without the ‘Combined Habitat Strategy’, over half of the CUs were predicted to have <50% probability of success (Fig. 3, Table S1). Further investment above $11.3M would benefit only three river-type sockeye CUs by increasing their probability of recovery to >50% at an extra $6M (Fig. 3).

### Sensitivity analysis

There was variation among the experts in the predicted costs and benefits of different strategies. Using the lower cost estimates, ‘Removal of Barriers to Fish Passage’ became the most cost-effective strategy (Table S2) and resulted in the same probability of achieving the conservation objective as ‘Limiting Future Industrial Development’ for less money ($69 900/year to remove five barriers over 20 years) under the 60% and 70% thresholds (Fig. S7). In the upper-cost scenario, the ‘Combined Habitat Strategy’ was 213% more expensive than the best-guess estimate, raising the average annual cost from $11M to $35M (Table S2), but was still selected in the complementarity analysis as it was the best strategy at improving the probability of success to >50% for 95% CUs (Fig. S7).

There was also wide uncertainty in the probability of each CU group achieving the objective under each strategy (Figs. S3 & S8). Experts predicted that under the worst-case scenario (i.e. experts’ lower benefit estimates), no CU would reach the >60% threshold under any strategy; the optimistic scenario (i.e., experts’ upper benefit estimates) suggested that most CUs would achieve a >60% probability of success if ‘Limiting Future Industrial Development’ or ‘Predation Control’ strategies were implemented (Fig. S8).

## Discussion

The PTM framework we implemented for BC’s Central Coast is a decision-support tool that can help guide conservation investment to maximize the benefit to Pacific salmon. We integrated Indigenous local knowledge into the analysis, through the design of the objectives and strategies, as well as their expert knowledge on the cost, feasibility and benefits, which we believe produced more relevant, inclusive, and legitimate results (Ban et al., 2018). This process relied on existing relationships among First Nations, and benefited from coordinated efforts led by the Pacific Salmon Foundation and CCIRA. Our study also demonstrated that the PTM framework can be applied not only to groups of threatened taxa or species, but also can include all levels of threat, and populations and ecotypes within species.

### Strategic planning in an era of high stakes for salmon

The strategic planning process revealed the urgency for conservation strategies and actions to support the recovery of Pacific salmon in Canada. Our analysis suggested a ‘business-as-usual’ baseline scenario will be insufficient to recover or maintain thriving salmon populations in BC’s Central Coast. Three quarters of CUs were estimated to have <50% chance of achieving this objective if no additional strategies are implemented (Table S1). However, if resources are allocated in a strategic manner our analyses suggest there is substantial scope to improve the overall status of salmon on BC’s Central Coast. In addition to the predicted benefits to salmon populations, the social, economic, and ecological benefits of implementing these proposed strategies potentially include: job provision, secure and stable fishing opportunities, health and well-being for affected communities, increased habitat protection, and greater opportunities for First Nation stewardship and governance.

The choice of decision-support framework is important because not all prioritization methods produce results that are equally cost-effective and beneficial (Giakoumi et al., 2013). If the goal is to conserve the most CUs per dollar spent, the associated costs, benefits, and feasibility of strategies should be compared across multiple species and CUs in a complementarity exercise, to avoid inefficient spending and maximize conservation outcomes (Martin et al., 2018). PTM has the potential to be a useful approach to guide the strategic and transparent allocation of new funding to maximize the return on investment for Pacific salmon.

At the regional level, PTM could serve as a model for agencies or First Nations interested in strategic planning and at the national level, it could inform the implementation of existing legislation and policies, such as the Canadian Species at Risk Act and the Canadian Wild Salmon Policy; both under restrictive budgets. There also is scope to explicitly include Indigenous social and cultural values of salmon in future PTM exercises by weighting different CU groups according to their relative importance to First Nation communities. This would provide a mechanism for ensuring that resources are preferentially directed to those CUs tied to First Nations cultures and communities.

### Cost-effective strategies for salmon conservation

The choice of strategy to maximize gains in salmon recovery on the Central Coast ultimately depends on the resources available (Fig. 3). Our analyses suggest that there are a few relatively inexpensive strategies that could provide immediate – though marginal – benefits to many salmon CUs, regardless of their biological status. ‘Limiting Future Industrial Development’ in critical spawning, rearing, and migration habitats was identified as a cost-effective option to safeguard and recover CUs, for an additional $150,000/year over the next 20 years above existing funding (Figs 3 & S6). Regardless of the conservation strategy chosen, the cost of monitoring and status assessment of salmon CUs ($0.7M/year) needs to be added to any budget, as this was considered necessary by experts. At budgets <$2M/year, we consider it would be important for decision-makers to decide which target threshold probability to aim for (i.e., >50, >60, or >70%), as the most cost-effective strategy depended on the chosen threshold (Fig. 3). If a larger budget of $11.3M were available per year, the ‘Combined Habitat Strategies’ were predicted to be highly beneficial. These actions include: i) protecting and restoring habitat and hydrology from forestry impacts, ii) restoring stream habitat to increase egg and juvenile survival, and iii) removal of barriers that limit fish passage and migration (Table 2). While these habitat strategies also had the highest uncertainty regarding the extent of habitat restoration and barrier removal required - leading to highly variable costs - they consistently outperformed other strategies in the sensitivity analyses (Fig. S7). Future research could refine these cost estimates.

Several uncertainties and caveats should be considered if implementing these strategies. It is inherently difficult for experts to predict the benefit of strategies into the future while accounting for multiple threats and underlying variable marine conditions that may reduce survival (e.g. Malick & Cox, 2016; Peterman & Dorner, 2012). This uncertainty is compounded by the logistical need to group together CUs that may have distinct (or unknown) biological status, life histories or threats. Management strategy evaluation frameworks (i.e., simulation modelling to quantify the predicted ability of strategies to meet multiple objectives) could be used to estimate expected outcomes while accounting for pervasive uncertainty (Punt, Butterworth, de Moor, De Oliveira, & Haddon, 2014), though pursuits for more knowledge should not be an excuse for delays in action (Martin et al., 2012). Opportunity costs (i.e. foregone profits) were not included in the analysis; further investigation into the economic benefits of industrial development, social preferences and risks would provide a more holistic assessment of this strategy.

### Conclusion

The PTM framework is a systematic approach that can be used to quantify trade-offs between costs and benefits of a diverse suite of strategies for Pacific salmon. The estimated benefits resulting from cost-effective strategies, combined with the cultural and economic importance of salmon to First Nations in this region, underscore the importance of prioritizing conservation strategies for Pacific salmon and their habitats in a systematic way. The status of salmon CUs in the Central Coast are relatively good compared to other regions such as southern BC, Washington and Oregon. This allows for a wider array of conservation management options, including proactive strategies that aim to safeguard CUs from the future risk of development. Even in regions with higher risks of local extirpation – where proposed strategies may be more expensive, less feasible and may result in lower probabilities of success – the PTM framework could provide a structured and transparent method to identify the most cost-effective options.

## Authors’ contributions

JCW, KC, EH, TGM, LK, BC and JDR conceived the ideas, designed methodology. JCW, KC, EH, TGM, LK, BC, JH and JDR collected the data; JCW and LK analyzed the data; JCW led the project and writing of the manuscript. KC, EH, BC, MJB, CF, AF, JWM, MHHP & JDR participated as experts in the elicitation process. All authors contributed critically to drafts and gave final approval for publication.

## Acknowledgements

We thank CCIRA and Stewardships Office staff from the Heiltsuk, Nuxalk, Kitasoo/Xai’xais and Wuikinuxv Nations for their support. We thank the following individuals for providing valuable advice and feedback: P. Siwallace, M. Reid, D. Chan, D. Rolston, B. Edgar, D. Neasloss, D. Dobson, D. Stewart, and all experts. Thank you to the Salmon Watersheds Program staff at Pacific Salmon Foundation for use of the Pacific Salmon Explorer, K. Kellock for producing maps and E. Jones, J. Belzile, C. Stevenson and L. Chalifour for facilitating workshop discussions. We thank D. Braun, W. Atlas & J. Walkus for comments on the manuscript. Mitacs, the Central Coast Indigenous Resource Alliance, Pacific Salmon Foundation and the Tom Buell Endowment Fund funded this project. Icons were sourced from Microsoft Office Professional Plus 2019.

## Data accessibility

Anonymized data are available on the Simon Fraser University repository [link to be provided once accepted].

## References

Ban, N. C., Frid, A., Reid, M., Edgar, B., Shaw, D., & Siwallace, P. (2018). Incorporate Indigenous perspectives for impactful research and effective management. Nature Ecology and Evolution, 2(11), 1680–1683. doi: 10.1038/s41559-018-0706-0

Barnas, K. A., Katz, S. L., Hamm, D. E., Dias, M. C., & Jordan, C. E. (2015). Is habitat restoration targeting relevant ecological needs for endangered species? Using Pacific Salmon as a case study. Ecosphere, 6(July), 110. doi: https://doi.org/10.1890/ES14-00466.1

Beechie, T., Pess, G., Roni, P., & Giannico, G. (2008). Setting River Restoration Priorities: A Review of Approaches and a General Protocol for Identifying and Prioritizing Actions. North American Journal of Fisheries Management, 28(3), 891–905. doi: 10.1577/M06-174.1

Bernhardt, E. S., Palmer, M. A., Allan, J. D., Alexander, G., Barnas, K., Brooks, S., … Sudduth, E. (2005). Synthesizing U.S. river restoration efforts. Science, 308(5722), 636–637. doi: 10.1126/science.1109769

Campbell, S. K., & Butler, V. L. (2010). Archaeological evidence for resilience of Pacific northwest salmon populations and the socioecological system over the last ~7,500 years. Ecology and Society, 15(1), 17. Retrieved from http://www.ecologyandsociety.org/vol15/iss1/art17/

Cannon, A., & Yang, D. Y. (2006). Early storage and sedentism on the Pacific Northwest Coast: ancient DNA analysis of salmon remains from Namu, British Columbia. American Antiquity, 71(1), 123–140. doi: https://doi.org/10.2307/40035324

Carwardine, J, Wilson, K. A., Watts, M., Etter, A., Klein, C. J., & Possingham, H. P. (2008). Avoiding costly conservation mistakes: The importance of defining actions and costs in spatial priority setting. PLoS ONE, 3(7), e2586. doi: https://doi.org/10.1371/journal.pone.0002586

Carwardine, Josie, Martin, T. G., Firn, J., Ponce-Reyes, R., Nicol, S., Reeson, A., … Chadès, I. (2019). Priority Threat Management for biodiversity conservation: a handbook. Journal of Applied Ecology, 56(2), 481–490. doi: 10.1111/1365-2664.13268

Carwardine, Josie, O’Connor, T., Legge, S., Mackey, B., Possingham, H. P., Martin, T. G., … Martin, T. G. (2012). Prioritizing threat management for biodiversity conservation. Conservation Letters, 5(3), 196–204. doi: 10.1111/j.1755-263X.2012.00228.x

Chadés, I., Nicol, S., van Leeuwen, S., Walters, B., Firn, J., Reeson, A., … Carwardine, J. (2015). Benefits of integrating complementarity into priority threat management. Conservation Biology, 29(2), 525–536. doi: 10.1111/cobi.12413

Connors, K., Jones, E., Kellock, K., Hertz, E., Honka, L., & Belzile, J. (2018). BC Central Coast: a snapshot of salmon populations and their habitats. Technical Report. Vancouver, BC: Pacific Salmon Foundation.

Cultus Sockeye Recovery Team. (2009). National conservation strategy for Cultus Lake sockeye salmon (Oncorhynchus nerka). Can. Tech. Rep. Fish. Aquat. Sci., 2846, viii + 46 p.

Evans, M. C., Tulloch, A. I. T., Law, E. A., Raiter, K. G., Possingham, H. P., & Wilson, K. A. (2015). Clear consideration of costs, condition and conservation benefits yields better planning outcomes. Biological Conservation, 191(November), 716–727. doi: 10.1016/j.biocon.2015.08.023

Fisheries and Oceans Canada. (2005). Canada’s Policy for Conservation of Wild Pacific Salmon. Vancouver.

Fisheries and Oceans Canada. (2018). Wild Salmon Policy 2018 to 2022 Implementation Plan. Retrieved from https://www.pac.dfo-mpo.gc.ca/fm-gp/species-especes/salmon-saumon/wsp-pss/ip-pmo/intro-eng.html

Garibaldi, A., & Turner, N. (2004). Cultural keystone species: implications for ecological conservation and restoration. Ecology and Society, 9(3), 1–18. doi: papers://59F6652F-E3FF-4FF7-BE89-9A861C9AA38C/Paper/p2970

Giakoumi, S., Sini, M., Gerovasileiou, V., Mazor, T., Beher, J., Possingham, H. P., … Katsanevakis, S. (2013). Ecoregion-Based Conservation Planning in the Mediterranean: Dealing with Large-Scale Heterogeneity. PLoS ONE, 8(10). doi: 10.1371/journal.pone.0076449

Good, T. P., Beechie, T. J., McElhany, P., McClure, M. M., & Ruckelshaus, M. H. (2007). Recovery planning for Endangered Species Act-listed Pacific Salmon: using science to inform goals and strategies. Fisheries, 32(9), 426–440. doi: https://doi.org/10.1577/1548-8446(2007)32[426:RPFESL]2.0.CO;2

Governor’s Salmon Recovery Office. (2018). 2018 State of the salmon in watersheds. Retrieved from Washington State Recreation and Conservation Office website: https://stateofsalmon.wa.gov/

Gustafson, R. G., Waples, R. S., Myers, J. M., Weitkamp, L. A., Bryant, G. J., Johnson, O. W., & Hard, J. J. (2007). Pacific salmon extinctions: Quantifying lost and remaining diversity. Conservation Biology, 21(4), 1009–1020. doi: 10.1111/j.1523-1739.2007.00693.x

Hemming, V., Burgman, M. A., Hanea, A. M., McBride, M. F., & Wintle, B. C. (2018). A practical guide to structured expert elicitation using the IDEA protocol. Methods in Ecology and Evolution, 9(1), 169–180. doi: 10.1111/2041-210X.12857

Hoekstra, J. M., Bartz, K. K., Ruckelshaus, M. H., Moslemi, J. M., & Harms, T. K. (2007). Quantitative Threat Analysis for Management of an Imperiled Species: Chinook Salmon (Oncorhynchus Tshawytscha). Ecological Applications, 17(7), 2061–2073. doi: 10.1890/06-1637.1

Holt, C. A., Cass, A., Holtby, B., & Riddell, B. (2009). Indicators of status and benchmarks for Conservation Units in Canada’s Wild Salmon Policy. In Canadian Science Advisory Secretariat Research Document (Vol. 58). Nanaimo, Canada, Canada.

Iacona, G. D., Sutherland, W. J., Mappin, B., Adams, V. M., Armsworth, P. R., Coleshaw, T., … Possingham, H. P. (2018). Standardized reporting of the costs of management interventions for biodiversity conservation. Conservation Biology, 32(5), 979–988. doi:10.1111/cobi.13195

Kareiva, P., Marvier, M., & McClure, M. (2000). Recovery and management options for spring/summer chinook salmon in the Columbia River Basin. Science, 290(5493), 977–979. doi: 10.1126/science.290.5493.977

Kehoe, L. J., Lund, J., Chalifour, L., Asadian, Y., Balke, E., Boyd, S., … Martin, T. G. (n.d.). Prioritizing conservation actions in heavily urbanized biodiverse socio-ecological systems. Unpublished.

Levi, T., Darimont, C. T., MacDuffee, M., Mangel, M., Paquet, P., & Wilmers, C. C. (2012). Using grizzly bears to assess harvest-ecosystem tradeoffs in salmon fisheries. PLoS Biology, 10(4), e1001303. doi: 10.1371/journal.pbio.1001303

Malick, M. J., & Cox, S. P. (2016). Regional-scale declines in productivity of pink and chum salmon stocks in western North America. PLoS ONE, 11(1), 1–23. doi: 10.1371/journal.pone.0146009

Mantua, N. J., Taylor, N. G., Ruggerone, G. T., Myers, K. W., Preikshot, D., Augerot, X., … Walters, C. J. (2009). The Salmon MALBEC Project: A North Pacific-scale Study to Support Salmon Conservation Planning. In North Pacific Anadromous Fish Commission Bulletin No 1060. Seattle, Washington, USA.

Marine Planning Partnership Initiative. (2015). Central Coast Marine Plan. Marine Planning Partnership Initiative, Heiltsuk, Kitasoo/Xai’Xais, Nuxalk and Wuikinuxv Nations, Province of British Columbia.

Martin, T.G., Kehoe, L., Mantyka-Pringle, C., Chades, I., Wilson, S., Bloom, R. G., … Smith, P. A. (2018). Prioritizing recovery funding to maximize conservation of endangered species. Conservation Letters, 11(6), e12604. doi: https://doi.org/10.1111/conl.12604

Martin, T. G., Nally, S., Burbidge, A. A., Arnall, S., Garnett, S. T., Hayward, M. W., … Possingham, H. P. (2012). Acting fast helps avoid extinction. Conservation Letters, 5(4), 274–280. doi: 10.1111/j.1755-263X.2012.00239.x

Martin, T. G, Burgman, M. A., Fidler, F., Kuhnert, P. M., Low-Choy, S., McBride, M., & Mengersen, K. (2012). Eliciting expert knowledge in conservation science. Conservation Biology, 26(1), 29–38. doi: 10.1111/j.1523-1739.2011.01806.x

Nelitz, M., Murray, C., & Wieckowski, K. (2008). Returning salmon: Integrated planning and the Wild Salmon Policy in B.C. Vancouver: ESSA Technologies for the David Suzuki Foundation.

Ogden, A. D., Irvine, J. R., O’Brien, M., Komick, N., Brown, G., & Tompkins, A. (2014). Canadian commercial catches and escapements of Chinook and coho salmon separated into hatchery- and wild-origin fish. NPAFC Doc. 1531. Retrieved from https://npafc.org/wp-content/uploads/2017/08/1531Canada.pdf

Peterman, R. M., & Dorner, B. (2012). A widespread decrease in productivity of sockeye salmon (Oncorhynchus nerka) populations in western North America. Canadian Journal of Fisheries and Aquatic Sciences, 69(8), 1255–1260. doi: 10.1139/f2012-063

Price, M. H. H., English, K. K., Rosenberger, A. G., MacDuffee, M., & Reynolds, J. D. (2017). Canada’s Wild Salmon Policy: an assessment of conservation progress in British Columbia. Canadian Journal of Fisheries and Aquatic Sciences, 74(10), 1507–1518. doi:10.1139/cjfas-2017-0127

Punt, A. E., Butterworth, D. S., de Moor, C. L., De Oliveira, J. A. A., & Haddon, M. (2014). Management strategy evaluation: best practices. Fish and Fisheries, 17(2), 303–334. doi: 10.1111/faf.12104

Roni, P., Beechie, T. J., Bilby, R. E., Leonetti, F. E., Pollock, M. M., & Pess, G. R. (2002). A Review of Stream Restoration Techniques and a Hierarchical Strategy for Prioritizing Restoration in Pacific Northwest Watersheds. North American Journal of Fisheries Management, 22(1), 1–20. doi: 10.1577/1548-8675(2002)022<0001:AROSRT>2.0.CO;2

Ruckelshaus, M. H., Levin, P., Johnson, J. B., & Kareiva, P. M. (2002). The Pacific Salmon Wars: What Science Brings to the Challenge of Recovering Species. Annu. Rev. Ecol. Syst, 33, 665–706. doi: 10.1146/annurev.ecolsys.33.010802.150504

Sakinaw Sockeye Recovery Team. (2005). National recovery strategy for sockeye salmon (Oncorhynchus nerka), Sakinaw Lake population, in British Columbia. Ottawa: Recovery of Nationally Endangered Wildlife.

Schoen, E. R., Wipfli, M. S., Trammell, E. J., Rinella, D. J., Floyd, A. L., Grunblatt, J., … Witmer, F. D. W. (2017). Future of Pacific Salmon in the Face of Environmental Change: Lessons from One of the World’s Remaining Productive Salmon Regions. Fisheries, 42(10), 538–553. doi: 10.1080/03632415.2017.1374251

Walters, C., English, K., Korman, J., & Hilborn, R. (2019). The managed decline of British Columbia’s commercial salmon fishery. Marine Policy, 101(December 2018), 25–32. doi: 10.1016/j.marpol.2018.12.014

Waples, R. S. (1991). Pacific salmon, Oncorhynchus spp., and the definition of “species” under the Endangered Species Act. Marine Fisheries Review, 53(3), 11–22. doi: http://spo.nmfs.noaa.gov/mfr533/mfr5332.pdf

