## Supplementary Information for "Prioritizing conservation actions for Pacific salmon in Canada"

### **Supporting Information: Prioritizing conservation actions for Pacific salmon in Canada**

### **Appendix S1: Detailed methods**

**Priority Threat Management framework**

The Priority Threat Management framework has eight steps, outlined in Figure 2 (main text). The first three steps were conducted in May 2018 during a day-long workshop in Vancouver, British Columbia with First Nation representatives and the Pacific Salmon Foundation, and several follow up conference calls. During a two-day workshop in June 2018, a panel of 19 experts quantified the costs, feasibility, and benefits of the management strategies (steps 4 & 5). These people had expertise in salmon threats, the feasibility and costs of conservation options in the Central Coast and/or the ecological response of salmon to each action. Experts included representatives from CCIRA and the Heiltsuk, Nuxalk, Kitasoo/Xai’xais, and Wuikinuxv Nations, academics, and resource managers, scientists and practitioners from government and non-governmental organisations. Efforts were made to have a broad diversity of knowledge and backgrounds of the expert group, as this has been shown to increase the accuracy and reliability of estimates (Bolger & Wright, 2011).

1. **Define objectives, scope and timeframe**

In the context of the Wild Salmon Policy, the objective was analogous to maximizing the number of CUs in the green status zone, which reduces the need for conservation intervention and allows for fishing opportunities, including catches for First Nation Food, Social, and Ceremonial (FSC) purposes, and commercial and recreational fishing (Fisheries and Oceans Canada, 2005). However, the ‘green’ status definition is complicated as status varies depending on the assessment metrics (e.g. stock recruitment analysis or historic spawners). We used the term thriving as it was also easier to understand and conceptualize by non-scientific experts and First Nation communities.

2. **Identify salmon CUs to conserve**

The ‘biodiversity features’ included in PTM exercises are usually threatened species or populations of concern, for which data are available. The 79 CUs were split into species groups. The sockeye CU group was further divided based on whether they are coastal or inland CUs, lake-type or river-type and their biological status (green, amber, red or data deficient based on percentiles of historic spawner abundance, as described in Connors et al. (2018)). The South Atnarko sockeye CU remained as a separate CU group given its poor conservation status (B. M. Connors & Atnarko Sockeye Recovery Planning Committee, 2016). This resulted in nine CU groups (Table 1).

3. **Identify threats, strategies and actions**

Strategies were initially proposed based on a review of existing literature, recovery plans, regional landscape-scale marine and watershed plans, habitat assessments, and preliminary workshops and meetings with experts (Connors et al., 2018; Marine Planning Partnership Initiative, 2015; Nelitz, Wieckowski, & Porter, 2007; Stalberg, Lauzier, Macisaac, Porter, & Murray, 2009). These strategies were then iteratively refined based on feedback from workshop participants resulting in ten strategies and associated actions (Table 2, Appendix S2).

Actions to reduce climate change and its effects on freshwater and marine habitats were not explicitly considered in this study given their spatial and governance scales were beyond the control of local resource managers.

Several actions regarding governance, management planning, and the development of fishing targets were included in an ‘Overarching harvest strategy’ that would support both the ‘Sustainable Commercial Harvest’ and ‘Sustainable Recreational Harvest’ strategies (Appendix S2).

4. **Estimate the costs and feasibility of actions**

The costs per action were divided into initial, annual, regular, and one-off costs (Carwardine et al., 2018). These costs were summed for each strategy and presented as the total present value (PV) over 20 years using a 4% discount rate (following past PTM exercises for public projects in Canada, Martin et al., 2018), and an average annual present value (AAPV). The costs of the enabling strategy of ‘Monitoring and Assessment’ were estimated but excluded from the cost-effectiveness analyses due to their limited ‘direct’ benefit to salmon. However, the experts considered monitoring and assessment an additional investment necessary to implement any other strategy, by tracking the status of populations (Price, English, Rosenberger, MacDuffee, & Reynolds, 2017). The costs of the ‘Overarching harvest strategy’ were divided between the ‘Sustainable Commercial Harvest’ and ‘Sustainable Recreational Harvest’ strategies. Best estimates of costs for each action were used in the analysis, but minimum and maximum costs were used in sensitivity analyses.

We did not estimate opportunity costs as a result of conservation strategies related to harvest (e.g. reduction in profits in the commercial and recreational fishing sectors due to changes in fishing gear, spatial closures, or restrictions in fishing licenses). Our rationale is that the future viability and profitability of the Pacific salmon industry depends on thriving populations of salmon; therefore, any short-term economic losses due to the implementation of a given strategy are expected to be offset by long-terms gains. We did not estimate the opportunity costs of limiting future industrial development, as this was outside the expertise of the expert group, though could be considered in future assessments.

The largest variation in costs came from uncertainty in the number of streams that required restoration of riparian vegetation, the number of roads to decommission to restore watershed hydrology, and the number and accessibility of artificial barriers to remove for fish passage.

5. **Identify benefits of strategies**

At the workshop, experts were given information from the online Pacific Salmon Explorer on current estimates of biological status for each CU, the extent and intensity of existing pressures on freshwater salmon habitat, past and current harvest rates, and anticipated industrial development projects in the region (Connors et al., 2018). Current biological status was calculated using recent spawner abundance relative to biological benchmarks derived from both percentiles of historic spawner abundance and stock-recruit analyses. Table 1 presents the current status of CUs using the historic spawner abundance benchmark, as this often produces the most precautious estimate of status (i.e. more often red or amber rather than green).

We asked experts to consider the effect of climate change and other future and emerging threats in their baseline estimates of probability of being thriving (or having green status) in 20 years. The baseline scenario also included all current management actions and policies that would presumably continue for the next 20 years. This included Integrated Fisheries Management Plans that outline annual harvest plans and management measures, and ongoing monitoring and assessment programs for salmon CUs. Each participant was given an Excel spreadsheet template to record their estimates of benefits. We used the modified Delphi method whereby experts provided their initial benefit estimates, and then had the chance to adjust their estimates after seeing box plots summarizing anonymous estimates by other experts (Mukherjee et al., 2015).

6. **Cost-effectiveness calculation**

The benefit metric per strategy per CU group used in the complementarity analysis (*Q_ij_*) was the probability of achieving a thriving population per strategy, accounting for the feasibility (*F_i_*), and calculated as the raw benefit of each strategy per CU group (*B_ij_*), and multiplied by the feasibility, which was then summed with the baseline probability (*B_0j_*, i.e. *P_0jk_* averaged across experts) (Chadés et al., 2015):

$$Q_{ij}= B_{ij} F_{i}+ B_{0j}$$

(Eqn S1)

The complementarity analysis was conducted following Chadés et al. (2015) and Kehoe et al. (n.d.).

For the sensitivity analysis, if a range of costs for each action were recorded, the upper and lower estimates also were tested in the simple ranking and complementarity analyses, using the ‘best-guess’ benefits scenario. Upper and lower limits of the experts’ estimates for the benefits were standardized to 80% confidence (Hemming, Burgman, Hanea, McBride, & Wintle, 2018), and then used in the analysis as the worst- and best-case scenarios, with the best-guess cost estimates.

**References**

Bolger, F., & Wright, G. (2011). Improving the Delphi process: Lessons from social psychological research. *Technological Forecasting and Social Change*, *78*(9), 1500–1513. doi: 10.1016/j.techfore.2011.07.007

Carwardine, J., Martin, T. G., Firn, J., Ponce-Reyes, R., Nicol, S., Reeson, A., … Chadès, I. (2019). Priority Threat Management for biodiversity conservation: a handbook. *Journal of Applied Ecology*, *56*(2), 481–490. doi: 10.1111/1365-2664.13268

Chadés, I., Nicol, S., van Leeuwen, S., Walters, B., Firn, J., Reeson, A., … Carwardine, J. (2015). Benefits of integrating complementarity into priority threat management. *Conservation Biology*, *29*(2), 525–536. doi: 10.1111/cobi.12413

Connors, B. M., & Atnarko Sockeye Recovery Planning Committee. (2016). *Atnarko Sockeye Recovery Plan*. Vancouver, BC: ESSA Technologies Ltd. for Nuxalk Nation.

Connors, K., Jones, E., Kellock, K., Hertz, E., Honka, L., & Belzile, J. (2018). *BC Central Coast: a snapshot of salmon populations and their habitats. Technical Report*. Vancouver, BC: Pacific Salmon Foundation.

Fisheries and Oceans Canada. (2005). *Canada’s Policy for Conservation of Wild Pacific Salmon*. Vancouver.

Hemming, V., Burgman, M. A., Hanea, A. M., McBride, M. F., & Wintle, B. C. (2018). A practical guide to structured expert elicitation using the IDEA protocol. *Methods in Ecology and Evolution*, *9*(1), 169–180. doi: 10.1111/2041-210X.12857

Kehoe, L. J., Lund, J., Chalifour, L., Asadian, Y., Balke, E., Boyd, S., … Martin, T. G. (n.d.). Prioritizing conservation actions in heavily urbanized biodiverse socio-ecological systems. *Unpublished*.

Marine Planning Partnership Initiative. (2015). *Central Coast Marine Plan*. Marine Planning Partnership Initiative, Heiltsuk, Kitasoo/Xai’Xais, Nuxalk and Wuikinuxv Nations, Province of British Columbia.

Martin, T. G., Kehoe, L., Mantyka-Pringle, C., Chades, I., Wilson, S., Bloom, R. G., … Smith, P. A. (2018). Prioritizing recovery funding to maximize conservation of endangered species. *Conservation Letters*, *11*(6), e12604. doi: https://doi.org/10.1111/conl.12604

Mukherjee, N., Hugé, J., Sutherland, W. J., McNeill, J., Van Opstal, M., Dahdouh-Guebas, F., … Koedam, N. (2015). The Delphi technique in ecology and biological conservation: applications and guidelines. *Methods in Ecology and Evolution*, *6*(9), 1097–1109. doi: 10.1111/2041-210X.12387

Nelitz, M., Wieckowski, K., & Porter, M. (2007). *Refining habitat indicators for strategy 2 of the wild salmon policy: Practical assessment of indicators*. Vancouver: Fisheries and Oceans Canada, prepared by ESSA Technologies.

Price, M. H. H., English, K. K., Rosenberger, A. G., MacDuffee, M., & Reynolds, J. D. (2017). Canada’s Wild Salmon Policy: an assessment of conservation progress in British Columbia. *Canadian Journal of Fisheries and Aquatic Sciences*, *74*(10), 1507–1518. doi: 10.1139/cjfas-2017-0127

Stalberg, H. C., Lauzier, R. B., Macisaac, E. A., Porter, M., & Murray, C. (2009). Canada’s Policy for Conservation of Wild Pacific Salmon: Stream, Lake, and Estuarine Habitat Indicators. *Canadian Manuscript Report of Fisheries and Aquatic Sciences*, *2859*, 1–151.

### **Appendix S2: Summary of strategies and actions for Central Coast Pacific salmon conservation**

**Overall objective** of the strategic planning exercise: Maximize the probability of achieving a healthy and thriving populations of Pacific salmon Conservation Units (CUs) on the Central Coast of British Columbia within the next 20 years (i.e., reduce the number of red-status CUs, and increase the number of green-status CUs, in the context of Canada’s Wild Salmon Policy). We define a healthy and thriving population as one that fulfills its ecological role and provides livelihood opportunities for present and future generations.

#### Enabling strategy: monitoring and assessment

The following actions were costed but were not included in the analyses, as they are an additional investment necessary for the implementation of any other strategy. We will assume the benefits to be the same across all strategies.

Actions within strategy

- Conduct integrated status assessments for CUs
- Conduct intensive monitoring of adult escapement and smolt abundance for CU indicator streams
- Establish monitoring programs for data deficient CUs

#### Overarching harvest strategy

The following actions were be costed and split across Strategies 1 (Commercial harvest) and Strategies 2 (Recreational harvest). We will assumed the benefits to be the same across these strategies.

Actions within strategy

- Establish quantitative management targets for groups of Conservation Units
- Establish co-management of fisheries between First Nations and Department of Fisheries and Oceans Canada regarding commercial and recreational catch limits, monitoring and enforcement. Co-develop Integrated Fisheries Management Plan
- Support and collaborate with First Nation Food, Social, and Ceremonial (FSC) initiatives to sustainably manage fisheries and improve FSC catch data

#### Strategy 1: Sustainable commercial harvest

Threat: **Commercial fisheries contributing to overfishing (i.e., harvest rates that exceed those predicted to maximize yield or that impair prospects for recovery) through directed catch, inception fisheries or by-catch.**

Goal: **Manage commercial harvest rates and catches with respect to management reference points to ensure CUs remain in green status, while rebuilding other CUs to green status over time.**

Actions within strategy

- Shift towards terminal fisheries, and limit mixed-stock catches to a lower percentage of total catch
- Improve enforcement of commercial fishery regulations
- Improve how the Pacific Salmon Treaty manages mixed-stock fisheries in Alaska to reduce impact on Central Coast CUs
- Reduce bycatch of non-target salmon species
- Research: routinely collect genetic samples for CUs to identify catch composition of mixed-stock fisheries

#### Strategy 2: Sustainable recreational harvest

Threat: **Recreational and sport fisheries contributing to overfishing (i.e., harvest rates that exceed those predicted to maximize yield or that impair prospects for recovery) through directed catch or catch-and-release methods.**

Goal: **Manage recreational harvest rates and catches with respect to management reference points to ensure CUs remain in green status, while rebuilding other CUs to green status over time.**

Actions within strategy

- Improve regulations to limit fleet and operations expansion by tourism and sport fishing operators; monitor/regulate independent sport fishers
- Monitor and restrict Fish and Wildlife permits
- Set more restrictive daily and annual harvest limits per person based on number of licenses and management reference points
- Set size limits for Chinook
- Limit excessive catch and release mortality by encouraging retention of legal-sized fish which count towards individual harvest limits (e.g. large numbers of coho should not be landed and released to enable Chinook harvest)
- Improve recreational catch monitoring and estimated mortality rates from catch and release fisheries for coho and Chinook
- Research: Estimate carrying capacity of tourism operators on the Central Coast, and their impact on salmon

#### Strategy 3: Watershed hydrology protection and water management

Threat: **Changes to watershed hydrology, water quality and quantity, and timing of flow in spawning and rearing habitat due to forestry and climate change. This includes floods or low water flows, and poor water quality due to sedimentation and runoff.**

##### **Goals:**

- **Maintain or restore the physical processes affecting natural basin hydrology, flow regime, and water quality and quantity**
- **Manage water use to optimize stream flows, water chemistry and sedimentation for salmonid spawning, incubation, rearing, and adult migration**
- **Maintain a complex river system that is less vulnerable to climate change.**

##### Actions within strategy

- Increase compliance, auditing and enforcement of logging regulations and forestry practices
- Identify streams vulnerable to high stream temperatures and variable flows. Then, implement relevant actions to ensure future sufficient flows
- Restore watershed vegetation to pre-logging conditions or other disturbance above existing work (e.g., reduce surface erosion from roads in priority areas by road decommissioning)

#### Strategy 4: Freshwater stream habitat restoration

Threat**: Loss or degradation of spawning and rearing habitat in streams and lakes (e.g., loss of big logs, woody debris, erosion of stream banks).**

Goal**: Maintain high quality fish and riparian habitat and restore spawning, rearing and migration habitats.**

##### Actions within strategy

- Conduct fine-scale habitat condition monitoring in priority areas using established assessment procedures (e.g., Watershed Assessment Procedure)
- Maintain and restore riparian habitat characteristics and processes (e.g., install large woody debris and engineering stream banks to ensure habitat complexity)
- Develop central database for habitat restoration actions, with measures of effectiveness

#### Strategy 5: Removal of barriers to fish passage and migration

Threat**: Culverts, roads, dams, etc. preventing adult and juvenile migration.**

Goal**: Remove existing artificial migration barriers to improve habitat connectivity.**

Actions within strategy

- Assess passability of barriers and stream crossings in Central Coast to identify barriers and identify priority barriers for removal
- Remove significant barriers (e.g., culverts) to upstream adult migration, or provide safe passage over these barriers (locations identified in first action) and remove significant barriers to juvenile dispersal to rearing habitats

#### Strategy 6: Marine and estuary habitat restoration and protection

Threat: **Loss and degradation of rearing habitat in estuaries (shipping, forestry practices, shellfish aquaculture, mining, pollution).**

Goal**: Maintain and restore high quality rearing estuary and marine habitats.**

##### **Actions in strategy**

- Research: Identify critical habitats for juveniles (i.e., nearshore rearing/benthic habitat) on BC’s Central Coast
- Protect eelgrass beds and other juvenile salmon rearing habitats - includes prohibiting dredging and dumping in nearshore habitats
- Restore estuaries degraded by forestry operations (e.g., marine loading sites, log sorts, helicopter-drop sites, booming areas, derelict vessels or fishing gear, abandoned sites)
- Implement strategies and best practices to reduce future impact from logging related activities in the estuaries and marine ecosystems
- Ensure that pollution policies and laws use international best practice guidelines and are implemented (i.e., develop and enforce provisions related to compensation for the destruction of fish and fish habitat)

#### Strategy 7: Limit impact of future industrial development in critical areas

Threat**: Potential loss or degradation of spawning and rearing habitat (freshwater and marine) from future mining, ports and harbors, aquaculture (other than salmon farming), refineries, pipelines, hydroelectric dams, windfarms, tourism (excluding sport fishing).**

**Salmon aquaculture is addressed in Strategy 10 and sport fishing is addressed in Strategy 2.**

Goal**: Reduce impacts of future industrial development on spawning, rearing and migration habitat.**

##### **Actions included in strategy**

- Restrict developments in estuaries (e.g., wind farms, aquaculture, and oil and gas infrastructure that will harm salmon) by developing protocol agreements between First Nations and proponents and maintaining existing tenures held by First Nations
- Manage water discharges from mines, gravel pits, and roads to reduce water pollution
- Research: Identify impacts of dumping sites (e.g., from clay mines) on out-migrating salmon
- Assess risks of shipping traffic to salmon

#### Strategy 8: Supplement small populations

Threat**: Small population sizes due to cumulative threats.**

Goal: **Use population enhancement if necessary to rebuild small or at-risk populations while carefully considering and reducing the negative impacts of hatcheries.**

##### Actions in strategy

- Evaluate current hatchery practices on Central Coast and conduct a risk analysis of their use and other options to supplement small salmon populations
- As a last resort for CUs in the critical red zone, initiate hatcheries. Mark all hatchery fish to ensure adequate monitoring and prohibit fishing on stocks enhanced for conservation
- Monitor and improve the effectiveness of hatcheries supplementing target CUs (e.g., parental based tagging)
- Provide full or partial fish ladders over natural barriers (e.g., waterfalls) affecting access to new spawning and rearing habitat (i.e., increase capacity of systems)
- Build spawning channels, where appropriate

#### Strategy 9: Predation control

Threat**: Predation from marine mammals and other predators.**

Goal**: Reduce impacts of predation by marine mammals (e.g., harbor seals and sea lions), and other predators.**

##### Actions included in strategy

- Identify importance and significance of predation by pinnipeds on Central Coast
- Conduct experimental culls (or traditional FN harvest) of pinnipeds to investigate effects at reducing predation pressure on juvenile and adult salmon
- Evaluate contribution of human-mediated predation and haul-outs (e.g., docks, seal restaurants)
- Trap sculpins when juveniles are migrating, especially for depressed populations
- Conduct hatchery releases at night to minimize predation risk by sculpins and trout
- Research: Understand historical relationship between FN and marine mammals/salmon

#### Strategy 10: Salmon aquaculture management

Threats**: Transmission of pathogens (parasites, bacteria and viruses) to wild salmon, water pollution and degradation of benthic habitat from salmon fish-farms.**

Goal**: Develop and implement a risk-adverse strategy to reduce the impacts of salmon aquaculture to wild salmon.**

Actions included in strategy

- Ensure existing salmon aquaculture facilities implement best practices that prevent the spread of parasites and disease to wild salmon
- Develop siting guidelines that preclude salmon aquaculture tenures to overlap with critical habitats for wild salmon and juvenile migratory routes
- Create tenures owned by First Nations to control new and existing aquaculture licenses
- Synthesize monitoring data to assess the impact of aquaculture on the Central Coast
- Assess potential interactions/risks to salmon populations transiting through aquaculture operations
- Incentivize land-based salmon aquaculture

#### Strategy 11: Combined harvest strategy

- Overarching harvest strategy
- Strategy 1: Sustainable commercial harvest management
- Strategy 2: Sustainable recreational harvest management

#### Strategy 12: Combined habitat strategy

- Strategy 3: Watershed hydrology protection and water management
- Strategy 4: Freshwater stream habitat protection and restoration
- Strategy 5: Remove artificial barriers to fish migration

#### Strategy 13: Combined supplementation and predation strategy

- Strategy 8: Supplement small populations
- Strategy 9: Predation control

#### Strategy 14: All strategies combined (excluding Monitoring and Assessment Enabling Strategy)

### Appendix S3: Supplementary Tables and Figures

##### Table S1: Probability of CUs within each CU group reaching a healthy and thriving status within 20 years under different strategies, while accounting for feasibility (*Q_ij_*, Eqn. S1). This metric was used as the ‘benefit’ in the complementarity analysis. BSL = baseline. DD = data deficient.

| CU group | Chinook | Chum | Coho | Pink | Coastal & Inland Lake-type Sockeye – Green | Coastal Lake-type Sockeye – Amber, Red & DD | Inland Lake-type Sockeye – Amber, Red & DD | South Atnarko Lake Sockeye | River-type Sockeye |
| --- | --- | --- | --- | --- | --- | --- | --- | --- | --- |
| BSL | 41.50 | 38.38 | 51.92 | 60.40 | 62.00 | 33.00 | 28.70 | 26.11 | 26.88 |
| S01 | 48.60 | 45.61 | 55.22 | 64.77 | 64.85 | 40.59 | 38.31 | 35.81 | 36.37 |
| S02 | 59.17 | 45.28 | 63.37 | 64.76 | 65.27 | 38.82 | 31.12 | 30.96 | 28.95 |
| S03 | 56.49 | 51.76 | 63.55 | 71.15 | 71.61 | 45.55 | 42.57 | 35.64 | 41.92 |
| S04 | 56.69 | 55.38 | 64.72 | 70.69 | 71.45 | 46.69 | 43.65 | 36.50 | 42.53 |
| S05 | 53.84 | 51.38 | 63.84 | 69.79 | 69.37 | 47.45 | 42.28 | 33.74 | 41.78 |
| S06 | 52.92 | 48.34 | 59.95 | 67.67 | 69.16 | 42.27 | 39.54 | 32.04 | 37.05 |
| S07 | 56.85 | 53.86 | 63.48 | 71.51 | 72.45 | 48.69 | 40.10 | 34.47 | 40.68 |
| S08 | 53.47 | 52.09 | 61.48 | 68.70 | 68.86 | 42.74 | 38.80 | 35.73 | 38.60 |
| S09 | 51.96 | 46.25 | 58.78 | 65.48 | 66.32 | 40.62 | 35.71 | 30.40 | 33.45 |
| S10 | 45.29 | 42.33 | 53.11 | 62.91 | 63.81 | 36.76 | 32.74 | 29.53 | 30.94 |
| S11 | 61.54 | 53.47 | 63.62 | 69.23 | 68.17 | 46.82 | 45.34 | 43.10 | 43.05 |
| S12 | 67.06 | 63.71 | 70.20 | 77.05 | 75.28 | 56.29 | 52.99 | 44.62 | 47.46 |
| S13 | 59.09 | 54.19 | 63.61 | 69.70 | 70.30 | 48.10 | 41.79 | 38.78 | 41.27 |
| All | 65.81 | 63.67 | 69.86 | 75.16 | 75.19 | 55.68 | 53.67 | 49.74 | 51.48 |
| No. of CUs/group | 6 | 9 | 6 | 5 | 8 | 35 | 6 | 1 | 3 |

##### Table S2: Sensitivity analysis using upper and lower estimates of costs per strategy; best estimates were used in main analysis.

|  |  | Average Annual Present Value (CAD millions) | | | Cost-effectiveness rank | | |
| --- | --- | --- | --- | --- | --- | --- | --- |
|  | Strategy name | Lower estimate | Best estimate | Upper estimate | Lower estimate | Best estimate | Upper estimate |
| S01 | Sustainable commercial harvest | 0.59 | 0.59 | 0.59 | 7 | 7 | 6 |
| S02 | Sustainable recreational harvest | 0.99 | 0.99 | 0.99 | 9 | 9 | 9 |
| S03 | Watershed protection | 0.18 | 0.54 | 0.89 | 3 | 5 | 4 |
| S04 | Stream restoration | 10.55 | 10.55 | 32.96 | 14 | 13 | 14 |
| S05 | Remove barriers to fish migration | 0.07 | 0.21 | 1.54 | 1 | 3 | 8 |
| S06 | Marine and estuary habitat restoration and protection | 0.79 | 0.79 | 0.79 | 6 | 6 | 5 |
| S07 | Limit future industrial development | 0.15 | 0.15 | 0.15 | 2 | 1 | 1 |
| S08 | Supplement small populations | 2.60 | 3.27 | 3.28 | 11 | 11 | 11 |
| S09 | Predation control | 0.11 | 0.11 | 0.11 | 4 | 2 | 2 |
| S10 | Aquaculture management | 0.09 | 0.09 | 0.09 | 5 | 4 | 3 |
| S11 | Strategy 1 + 2 | 1.59 | 1.59 | 1.59 | 8 | 8 | 7 |
| S12 | Strategy 3 + 4 + 5 | 10.81 | 11.30 | 35.39 | 12 | 12 | 12 |
| S13 | Strategy 8 + 9 | 2.72 | 3.39 | 3.39 | 10 | 10 | 10 |
| S14 | All strategies combined | 16.147 | 17.30 | 41.39 | 13 | 14 | 13 |

##### Figure S1: Maps of Conservation Units (CUs) for coho, river-type sockeye, and pink (even and odd year runs) within the Central Coast of British Columbia, Canada, with the study region outlined in green showing the traditional use territories of the Heiltsuk, Kitasoo/Xai’xais, Nuxalk, and Wuikinuxv Nations.

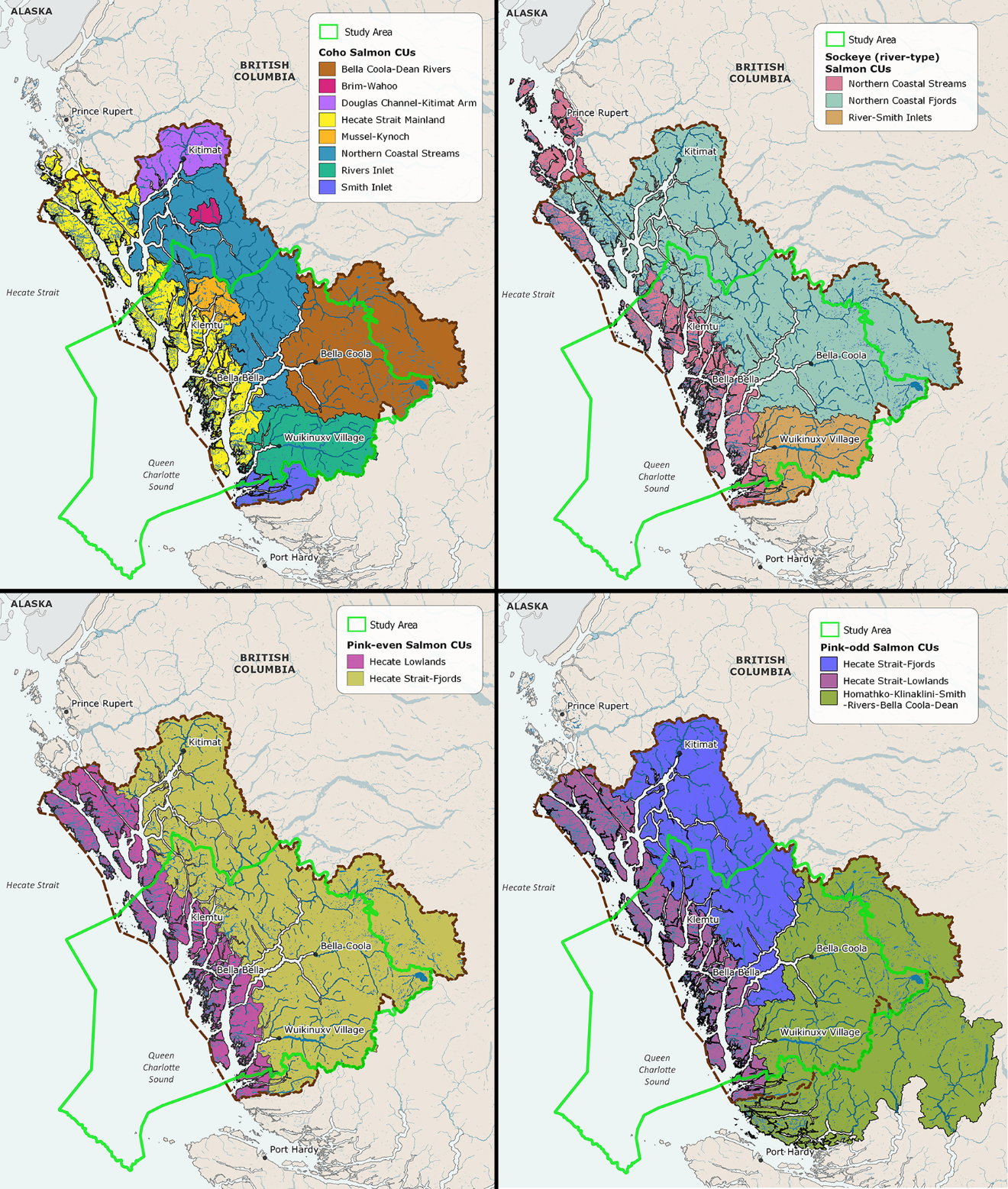

##### Figure S2: Lake-type sockeye salmon Conservation Units (CUs) within the Central Coast of British Columbia, Canada, with the study region outlined in green showing the traditional use territories of the Heiltsuk, Kitasoo/Xai’xais, Nuxalk, and Wuikinuxv Nations.

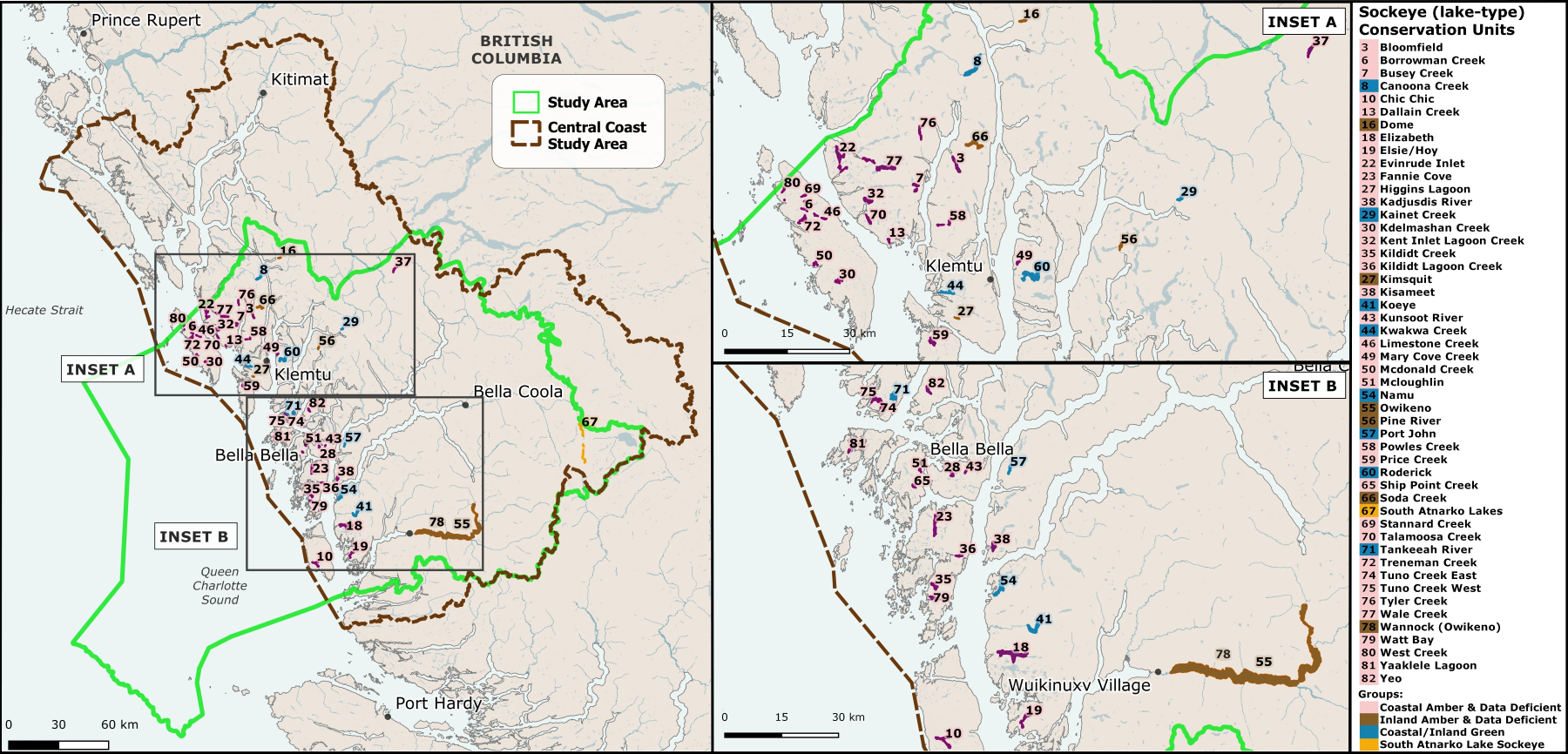

##### Figure S3: Box plots of the experts’ final best-guess estimates of the probability of each Conservation Unit (CU) group achieving the conservation objective (thriving populations) for each strategy within the next 20 years (i.e. *P_ijk_*). BSL = baseline scenario. Strategy numbers correspond to Table S2.

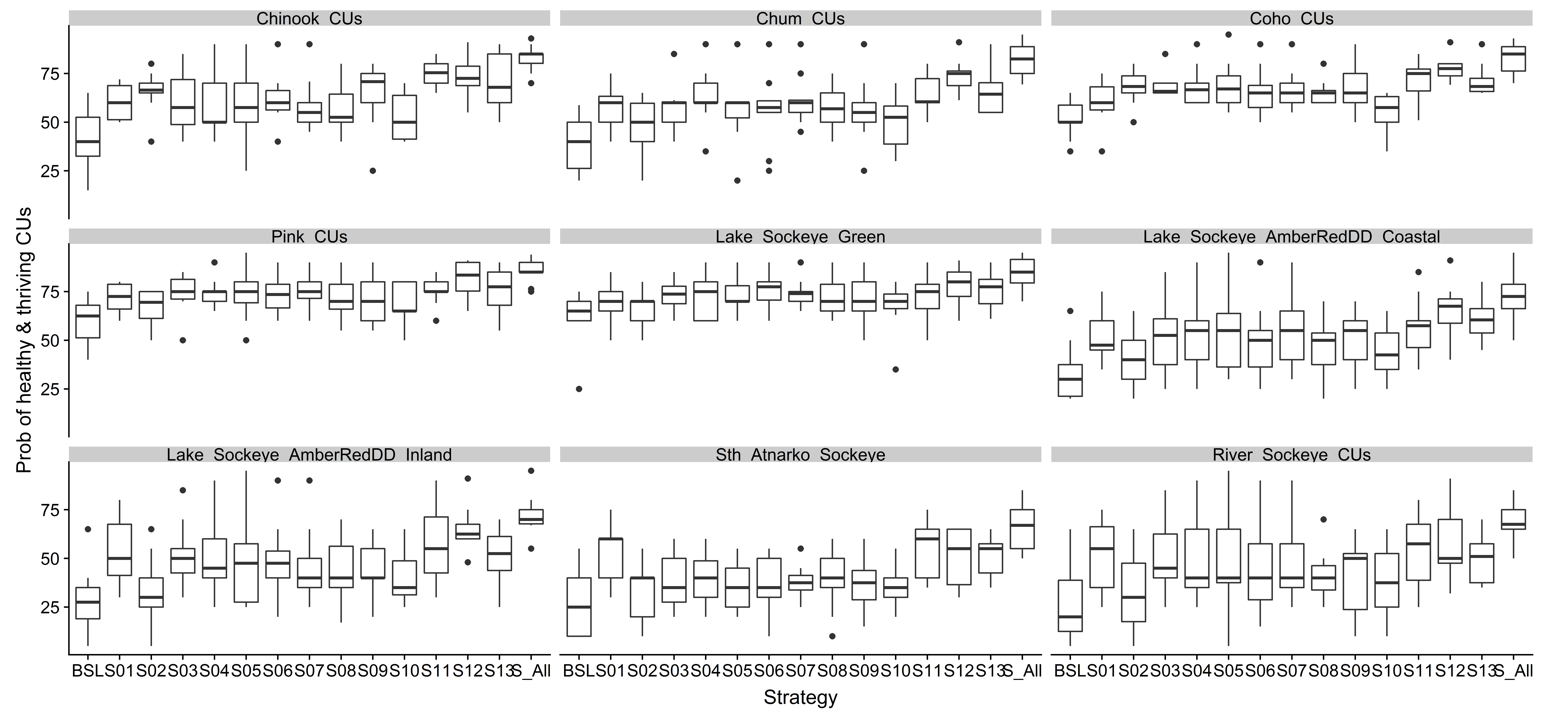

##### Figure S4: Cost-effectiveness ranking of strategies for Pacific salmon conservation on Central Coast British Columbia. Light bars are individual strategies and dark bars are combined strategies (blue = harvest strategies, green = habitat strategies, orange = other strategies, brown = all strategies combined).

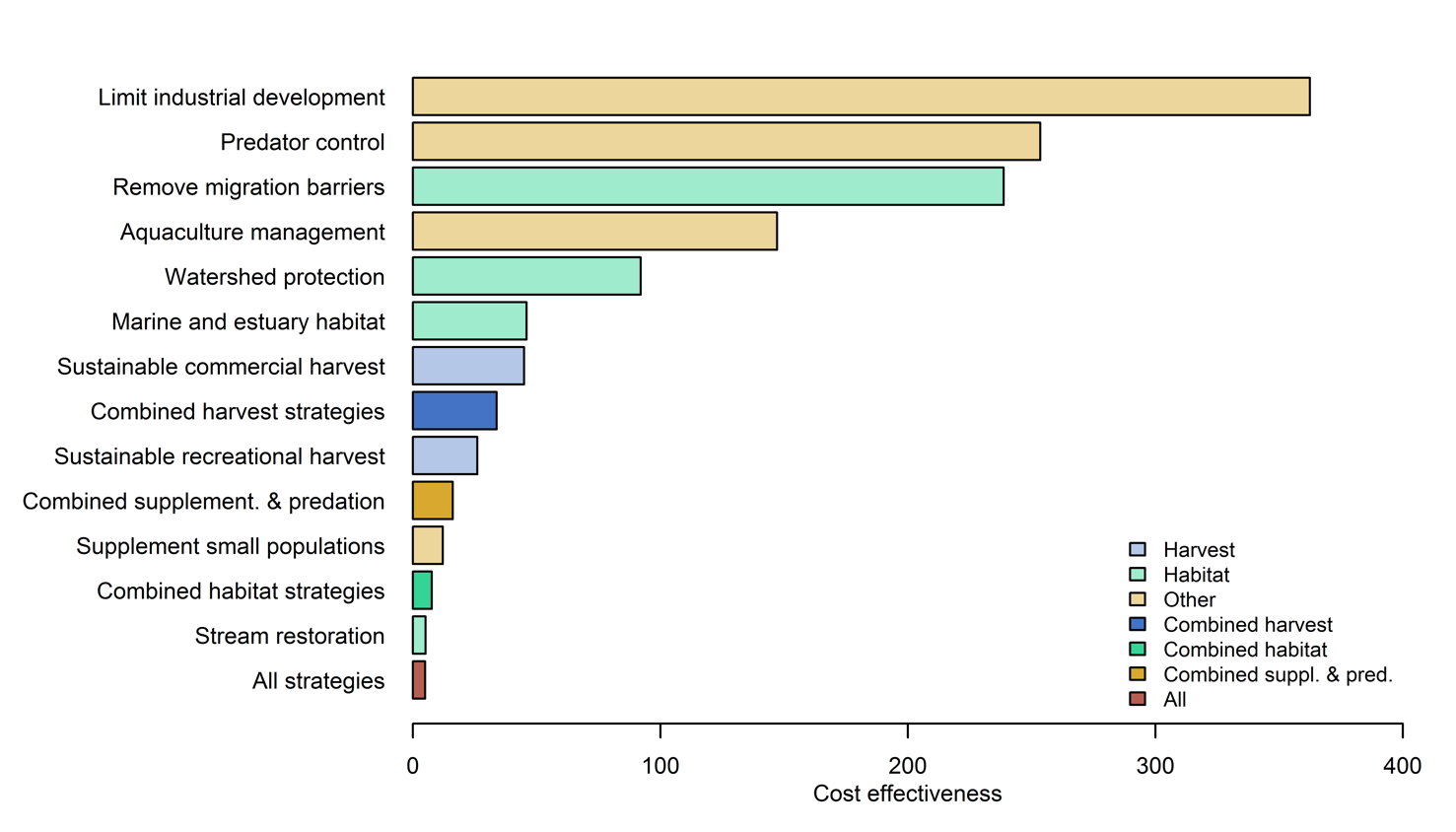

##### Figure S5: Benefits (*B_ij_*), feasibility, and cost of strategies for Pacific salmon conservation on the Central Coast British Columbia. Light bars are individual strategies and dark bars are combined strategies (blue = harvest strategies, green = habitat strategies, orange = other strategies, brown = all strategies). Benefits are the cumulative benefit of a strategy summed across all CU groups, weighted by the number of CUs in each group.

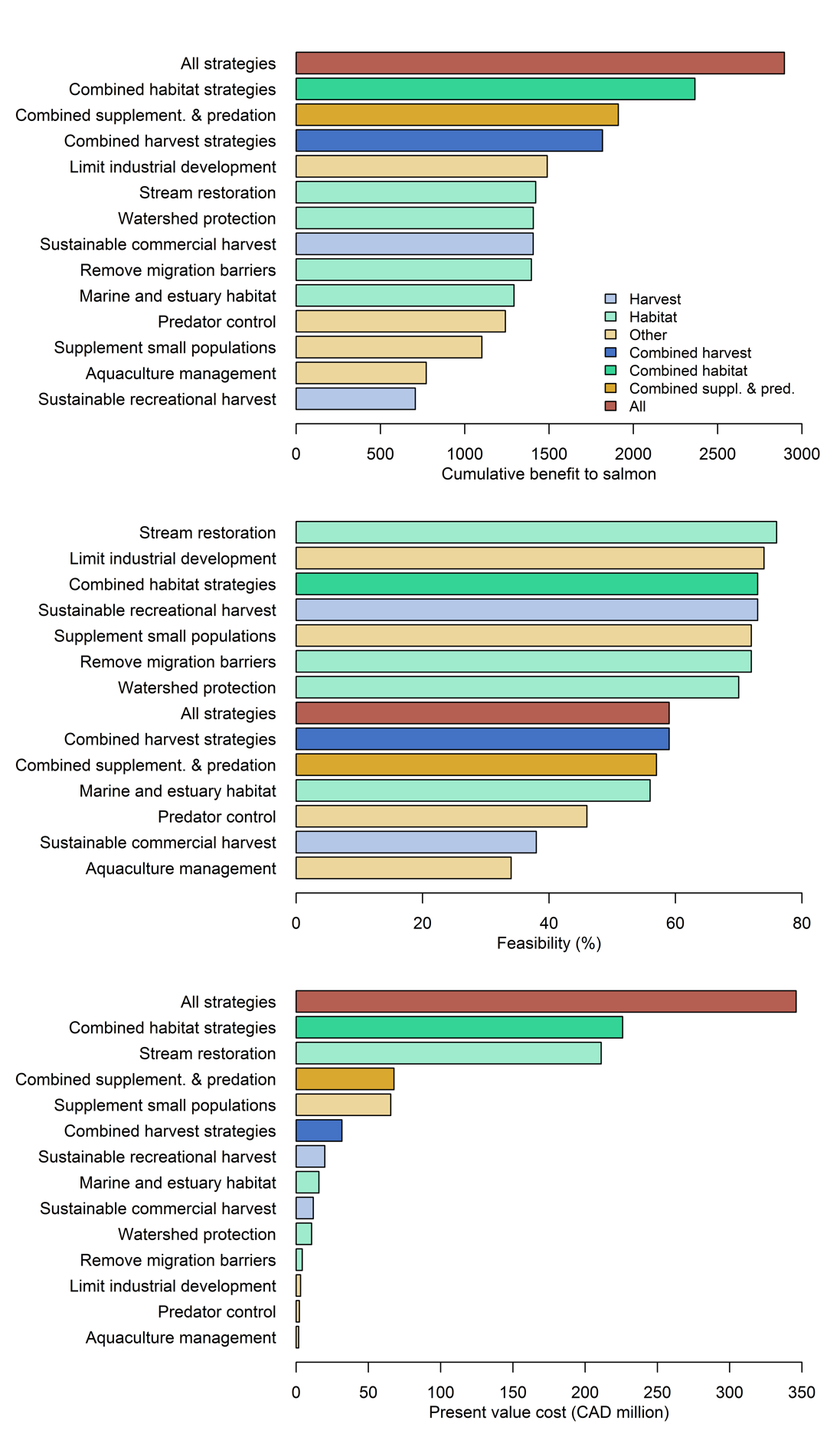

##### Figure S6: Benefits (*B_i_*, i.e., probability of achieving objective with strategy – probability of achieving objective without strategy) across all groups of Conservation Units (CUs). These benefits were weighted by the number of CUs per group and summed to calculate a cumulative benefit estimate per strategy shown in Fig. S5a (Eqns 1 & 3 main text).

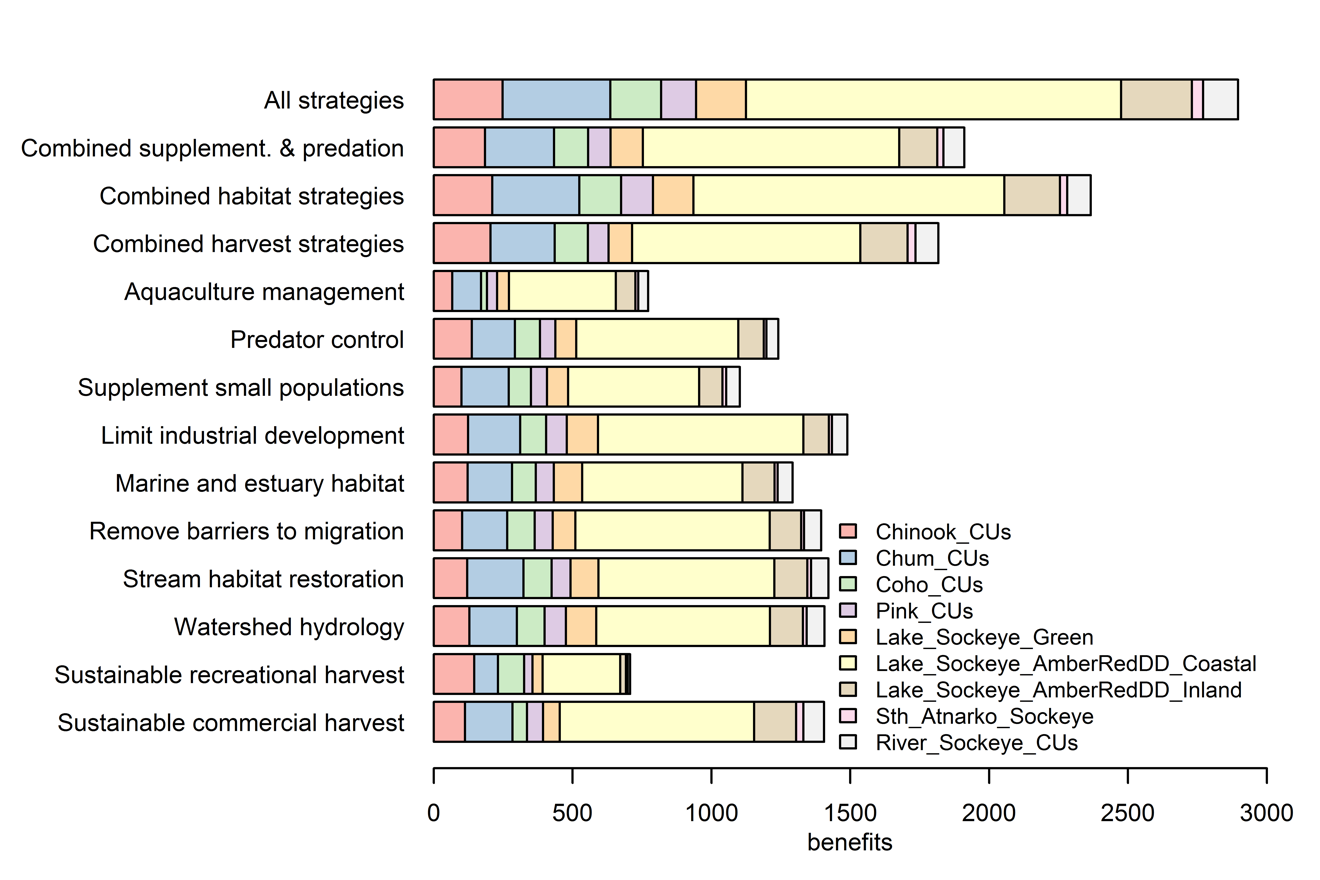

##### Figure S7: Results of cost sensitivity analysis of complementarity analysis with lower cost estimates (top plot), best-guess estimates (middle plot) and upper cost estimates (bottom plot).

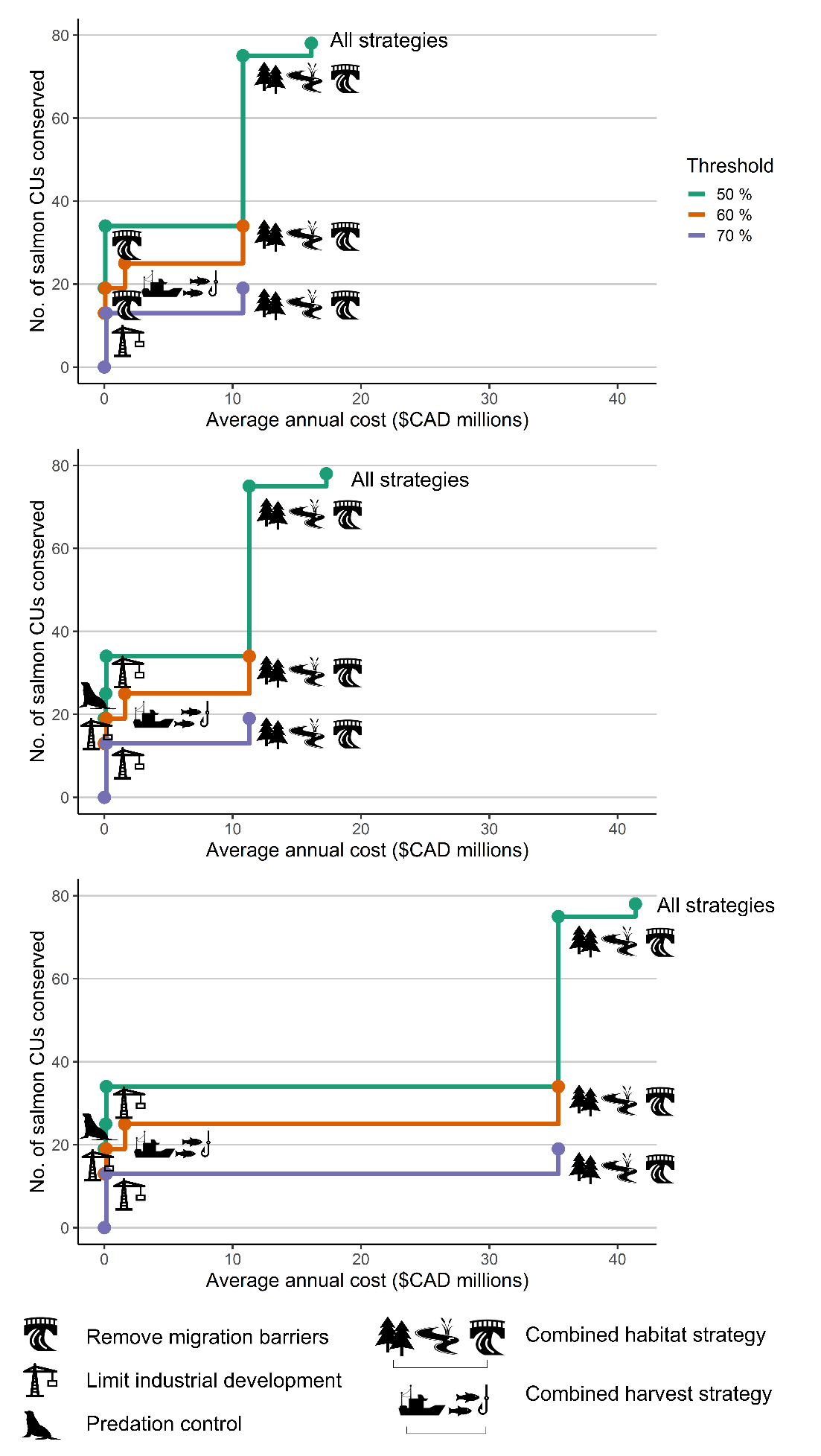

##### Figure S8: Benefit sensitivity analysis of complementarity analysis with lower (top plot), best-guess (middle), and upper (bottom plot) benefit estimates from experts (i.e., pessimistic, best-guess and optimistic scenarios).

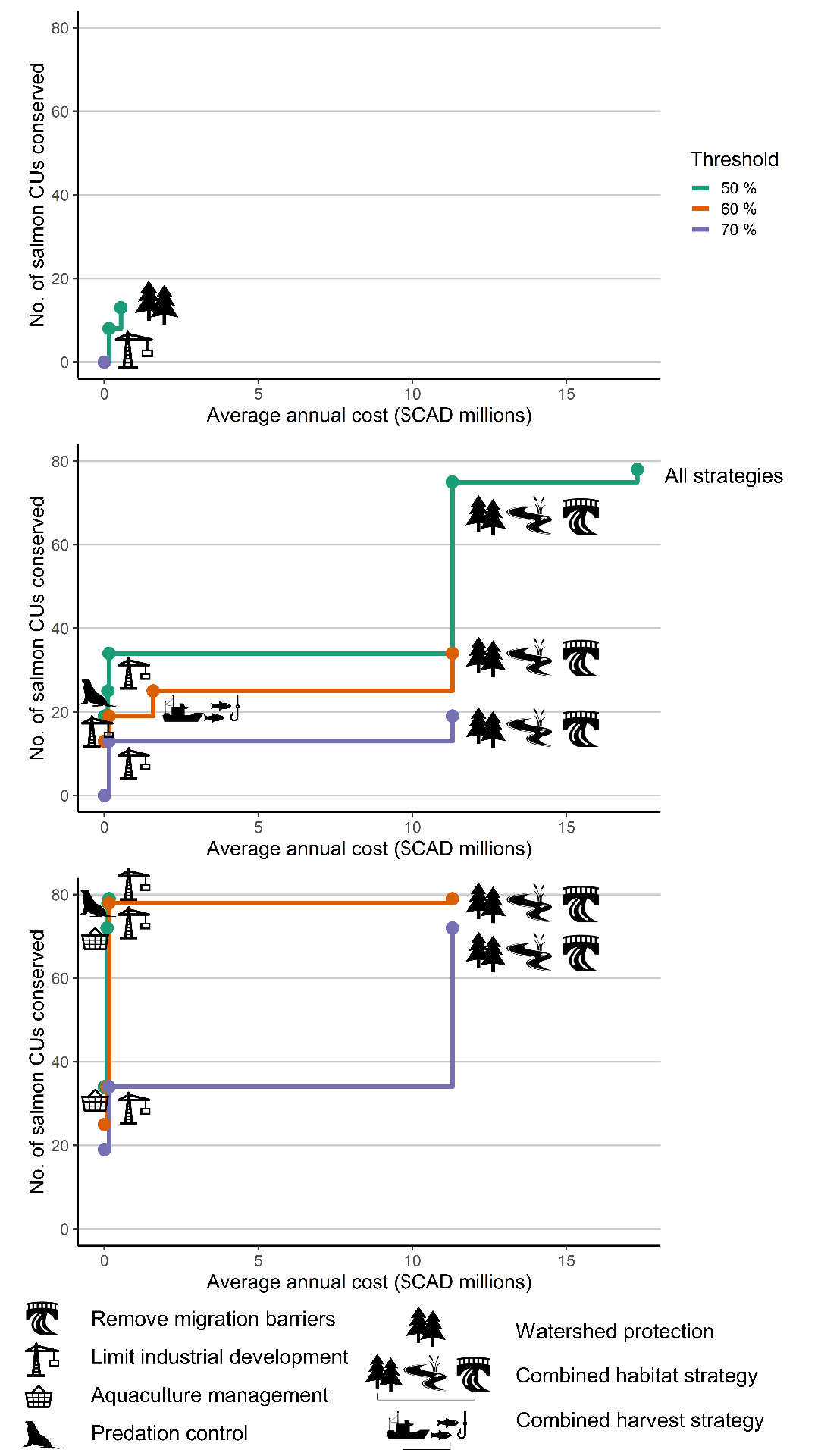
